# Postsynaptic receptors regulate presynaptic transmitter stability through trans-synaptic bridges

**DOI:** 10.1101/2022.09.10.507343

**Authors:** Swetha K. Godavarthi, Masaki Hiramoto, Yuri Ignatyev, Jacqueline B. Levin, Hui-quan Li, Marta Pratelli, Jennifer Borchardt, Cynthia Czajkowski, Laura N. Borodinsky, Lora Sweeney, Hollis T. Cline, Nicholas C. Spitzer

## Abstract

Stable matching of neurotransmitters with their receptors is fundamental to synapse function, to achieve reliable and robust communication in neural circuits. Presynaptic neurotransmitters regulate selection of postsynaptic transmitter receptors. However, whether postsynaptic receptors regulate selection of presynaptic transmitters is unknown. Here we show that blockade of postsynaptic acetylcholine receptors at the neuromuscular junction leads to loss of the cholinergic phenotype in motor neurons and stabilization of an earlier, developmentally transient glutamatergic phenotype. Exogenous postsynaptic expression of GABA_A_ receptors leads to the stabilization of an earlier, developmentally transient GABAergic motor neuron phenotype. Both acetylcholine receptors and GABA receptors are linked to presynaptic neurons through trans-synaptic bridges. Knock-down of different components of these trans-synaptic bridges prevents stabilization of the cholinergic and GABAergic phenotypes. We conclude that this bidirectional communication enforces a match between transmitter and receptor and ensures the fidelity of synaptic transmission. Our findings suggest a role of dysfunctional transmitter receptors in neurological disorders that involve the loss of the presynaptic transmitter.

## Main

Postsynaptic receptors at neuronal synapses are often viewed as passive responders to presynaptic neurotransmitters. For example, dendrites respond to uncaged glutamate or GABA by forming spines that express glutamate or GABA receptors^1,2^, filopodia respond to release of glutamate from developing axons, leading to physiological and morphological maturation^3^, and cultured glutamatergic neurons form functional glutamatergic synapses with skeletal muscle cells^4^. Also, when neurotransmitters switch, the postsynaptic cell responds to the newly expressed transmitter by expressing a matching receptor^5–8^. Activity-driven retrograde signaling by endocannabinoids and neurotrophins from the postsynapse regulates presynaptic neuron survival, dendrite formation and growth, and synaptogenesis^9,10^. Do changes in the receptors on postsynaptic cells alter transmitter expression in their presynaptic neurons? Here we demonstrate that retrograde signaling by postsynaptic transmitter receptors is necessary and sufficient to stabilize expression of their cognate transmitter in presynaptic neurons and that signaling is blocked by disruption of trans-synaptic bridges.

### Blocking endogenous acetylcholine receptors at the neuromuscular junction

We first tested whether blockade of acetylcholine receptors (AChR) of cholinergic neuromuscular junctions in *Xenopus* embryos affects the expression of acetylcholine in motor neurons. Agarose beads containing AChR antagonists, pancuronium or curare, were implanted into developing mesoderm at 1 day post fertilization (dpf) for local, diffusive drug delivery (**Fig. 1a**). Larvae were wholemount immunostained for choline acetyltransferase (ChAT), the enzyme that synthesizes ACh, and for synaptic vesicle protein 2 (SV2), a marker of nerve terminals, to determine the capacity of nerve terminals for ACh synthesis. At 2 dpf, the staining of terminals running along the myocommatal junctions at the boundaries between chevrons of myocytes was not different from the staining in saline bead controls (**Fig. 1b** and **1e**). By 3 dpf, the staining for ChAT had decreased to 19% of controls with no change in staining for SV2 (**Fig. 1c** and **1e**). By 4 dpf the staining for ChAT had fallen to 5% of controls (**Fig. 1d** and **1e**), while SV2 staining was 73% of controls (**Fig. 1e**; **Suppl Figs. 1, 2, 3 and Suppl Table 1**). These results indicate that the loss of ChAT precedes the loss of nerve terminals and suggest that blockade of postsynaptic AChR impairs the stability of ChAT expression. This finding is consistent with the observation of a decrease in the presynaptic active zone protein Bruchpilot, associated with presynaptic transmitter synthesis and release^11^, following blockade of postsynaptic receptor subunits^12^ that occurs early in homeostatic presynaptic scaling^13,14^. The decrease is reversed only when the receptor function is restored by expression and insertion of a different, reserve receptor subunit^14,15^. In the present case, the absence of reserve receptor subunits of the skeletal muscle ACh receptor^16^ facilitated identification of the postsynaptic receptor-dependence of presynaptic transmitter stability. Interestingly, the loss of AChR precedes the loss of nerve terminals in a mouse model of myasthenia gravis (MG)^17^. MG is often the result of an autoimmune attack on AChR function^18– 20^. Our findings suggest that reduced levels of presynaptic ACh, in addition to loss of AChR, contribute to the ensuing muscle fatigue that is also observed in congenital MG resulting from ChAT deficiency^19^.

**Fig 1.**
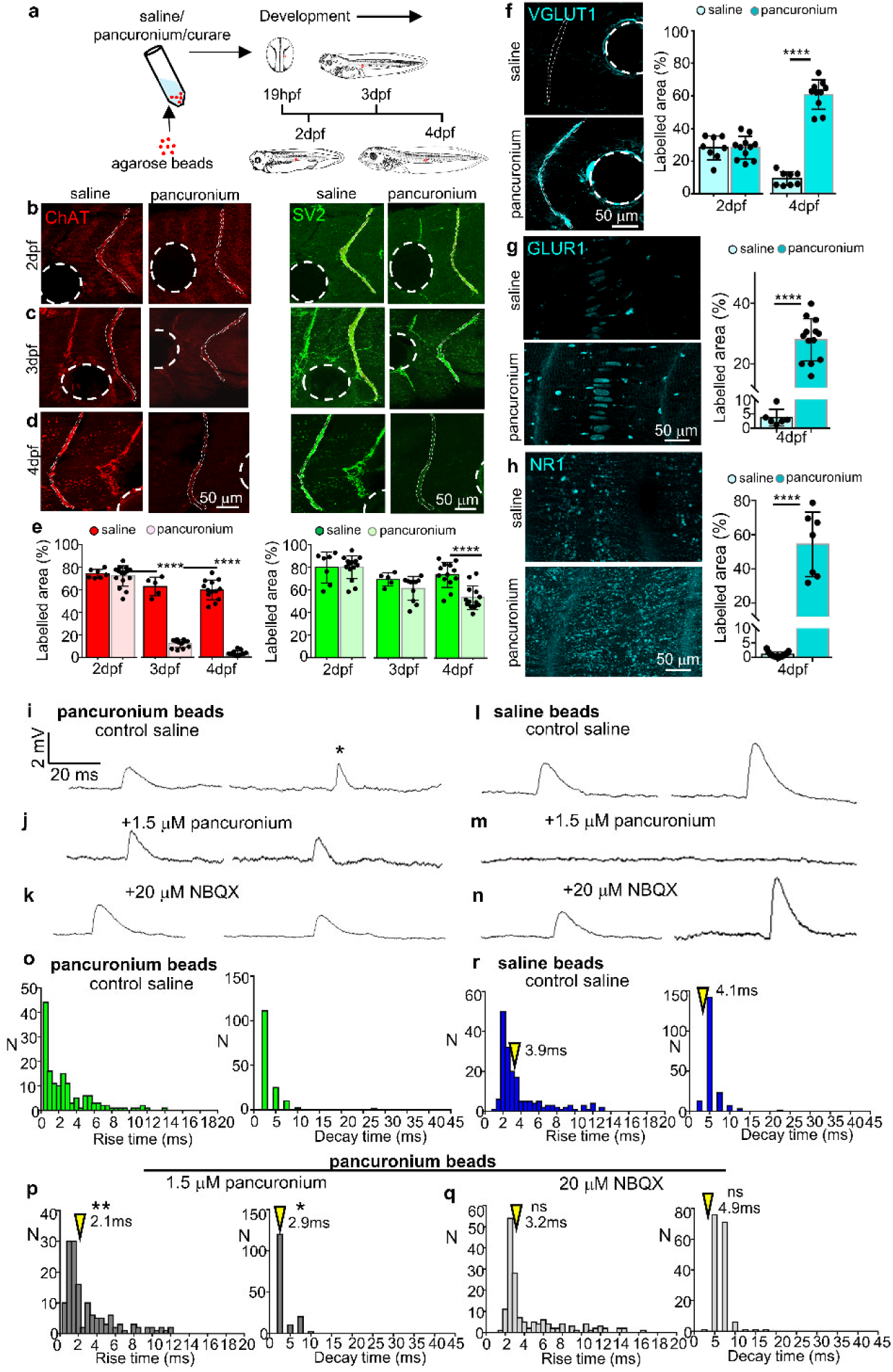
Local block of AChR in myocytes reduces ChAT expression in innervating axons and induces a glutamatergic phenotype. (**a**) Experimental design. A single agarose bead loaded with pancuronium, curare, or saline control was implanted into the *Xenopus* mesoderm at 19hpf. (**b-d**) Bead-implanted larvae were examined at 2dpf, 3dpf and 4dpf for ChAT and SV2. Dotted lines outline myocommatal junctions analyzed for quantification of staining area. Dashed lines indicate positions of beads. (**e**). Area of expression quantified for ChAT and SV2. N≥7 larvae. (**f-h**). Expression and quantification of VGLUT1, GLUR1 and NR1 in 4dpf myotome of control and pancuronium-loaded agarose bead-implanted larvae. N≥8 larvae. ****p<0.0001, unpaired two-tailed t-test. (**i-k**) Recordings from pancuronium-bead-implanted larvae reveal rapid rise and rapid decay AMPAR-mediated-PSP-like mEPPs (asterisk) that are pancuronium-resistant and NBQX-sensitive, as well as pancuronium-sensitive and NBQX-resistant mEPPs with rise and decay times similar to those described for nicotinic receptor-mediated-EPPs. (**l-n**) Recordings from saline-bead-implanted larvae reveal only pancuronium-sensitive mEPPs. (**o-r**) Rise and decay time distributions for mEPPs in myocytes of pancuronium-bead implanted larvae and saline-bead implanted larvae. N, number of mEPPs. >155 mEPPs (≥3 larvae, 4dpf) for each group. Only mEPPs with decay times fit by single exponentials were included. Resting potentials were held near −60 mV. Arrowheads indicate median values. Kolmogorov-Smirnov test compared rise time and decay time in (**r**) with respective rise and decay time in (**p**) and (**q**). *p<0.05, **p<0.01, ns not significant.

Normally, embryonic *Xenopus* motor neurons express glutamate in their cell bodies at 1 dpf and its level decreases by 2 dpf as ChAT begins to be detected^21^. By 3 dpf glutamate is no longer detected immunohistochemically. Blockade of AChR with pancuronium led to increased expression of a vesicular glutamate transporter (VGLUT1) in motor neuron terminals (**Fig. 1f**; **Suppl Fig. 4**), suggesting the stabilization of a glutamatergic phenotype. The expression of VGLUT1 was accompanied by a corresponding increase in expression of glutamatergic AMPA and NMDA receptor subunits in myocytes (**Fig. 1, g** and **h**). Intracellular recordings from these myocytes at 4 dpf yielded two classes of miniature endplate potentials (mEPPs), recognized by their rise and decay times (**Fig. 1, i** to **r**) and their frequency, with a mean frequency overall of 0.6±0.0 sec^-1^. Rapid rise-rapid decay mEPPs with a mean frequency of 0.4±0.0 sec^-1^ were blocked by NBQX, indicating that they depended on AMPA receptors. The mEPPs remaining after the NBQX block had slower rise and decay times typical of AChR and a mean frequency of 0.1±0.0 sec^-1^. The mEPPs recorded in the presence of NBQX were eliminated by the additional application of pancuronium. Control larvae implanted with a saline bead had a mean frequency of 1.0±0.1 sec^-1^. Because quantal content is proportional to mEPP frequency^22,23^, the higher frequency of glutamatergic mEPPs than cholinergic mEPPs implies larger evoked glutamate release in response to ACh receptor blockade that is a feature of homeostatic presynaptic scaling. The local block of AChR did not result in immunostaining for GABA (fig. S5), in agreement with the earlier reports of increase in the number of neurons expressing glutamate, but not GABA or glycine, that occurs following a reduction in neuronal activity^5,24^. These results show that local block of ACh receptors leads to stabilization of expression and function of another excitatory transmitter, glutamate.

### Expressing exogenous GABA receptors in embryonic myocytes

Embryonic *Xenopus* motor neurons express GABA as well as glutamate in their cell bodies and in their axons^21^ (**Suppl Fig. 6**). Ordinarily, the level of GABA decreases by 2 dpf as ChAT appears, and GABA has disappeared by 3 dpf. We expressed GABA_A_ receptors in a small number of embryonic myocytes^25^ to learn whether this would stabilize expression of GABA in the motor neurons that innervate them. We co-injected the transcripts for GABA_A_ receptor α1-EGFP, β2, and γ2 subunits into the ventral blastomeres (V) at the 4-cell stage, to achieve assembly of GABA_A_Rαβγ receptors. Injection of α1-EGFP transcripts alone served as control, since expression in this case is restricted to the cytoplasm and does not appear in the plasma membrane^26^ (**Suppl Fig. 7**). The GFP-tag on the α1 subunit labeled the transfected myocytes. Strikingly, at 3 dpf, expression of GABA_A_Rαβγ receptors in myocytes, but not GABA_A_Rα alone, led to expression of GABA and VGAT in the nerve terminals that innervate them (**Fig. 2; Suppl Fig. 8**). ChAT was coexpressed with GABA and VGAT, suggesting that the terminals are processes of motor neurons (**Suppl Fig. 9, a** to **h**). Further support that the nerve terminals expressing GABA and VGAT are from motor neurons came when we traced GABA-labeled axons in the myotome back to spinal cord cell bodies expressing motor neuron transcription factor Hb9 (data not shown). This conclusion was strengthened by observation that GABA and Hb9, as well as GAD67 and ChAT, are coexpressed in neuronal cell bodies (**Suppl Fig. 9, i** to **l**). Moreover, neurons expressing ChAT and GABA in their cell bodies expressed the Lim3 and not the rAldh1a2 transcription factor. This result demonstrated that the motor neurons were formed during the primary wave of neurogenesis and not by a population formed later (**Suppl Fig. 10**). Expression of GABA_A_Rαβγ stabilized the GABAergic phenotype but did not stabilize expression of VGLUT1 or glycine in innervating axons (**Suppl Fig. 11**).

**Fig 2.**
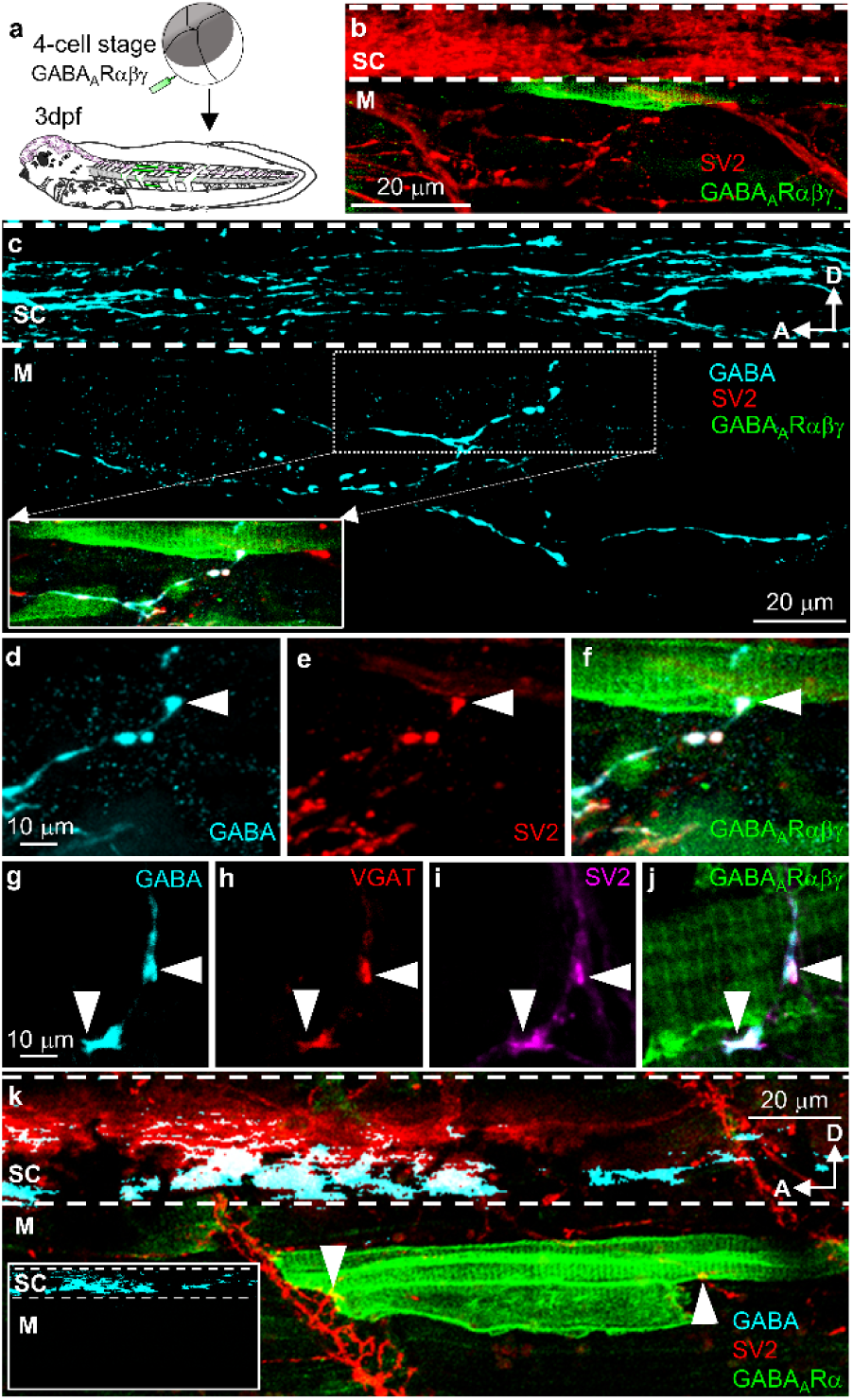
GABA_A_Rαβγ expression in myocytes leads to GABA expression in axons that innervate them. (**a**) Injection of ventral blastomeres (V) with GABA_A_Rαβγ mRNA at 4-cell stage results in myocyte-specific GABA_A_R expression. (**b**) Sparse expression of GABA_A_Rαβγ in the trunk myotome. SV2 labels neuronal axons in the spinal cord and in the trunk myotome. (**c**) In an expansion of the same field of view, staining for GABA reveals GABA+ axons in the spinal cord and coursing ventrally and posteriorly over the trunk myotome. *Inset:* A GABA+SV2+ axon contacts a GABA_A_Rαβγ-expressing GFP+ myocyte (box). (**d-f**) Higher magnification of contact in (**c**) (arrowheads). (**g-j**) A GABA+VGAT+SV2+ axon contacts a different GABA_A_Rαβγ myocyte. (**k**) Injection of ventral blastomeres (V) with GABA_A_Rα mRNA results in sparse expression of GABA_A_Rα in the trunk myotome. SV2+ axons contact a GABA_A_Rα-expressing GFP+ myocyte (arrowheads). *Inset*, GABA+ axons are restricted to the spinal cord. M, trunk myotome; SC, spinal cord; D, dorsal; A, anterior. 3dpf.

The expression of GABA in motor neuron axons at 2 dpf in these embryos was increased specifically in axons contacting myocytes expressing GABA_A_Rαβγ. Myocytes expressing GABA_A_Rα did not elicit axonal GABA expression (**Suppl Fig. 12**), suggesting that GABA was stabilized by the presence of the cognate postsynaptic receptor. The stabilization of GABA in axons innervating GABA_A_Rαβγ-expressing myocytes persisted up to 7 dpf (**Suppl Fig. 13**). To further test whether GABA_A_Rαβγ-expressing myocytes were specific in stabilizing the presynaptic GABAergic phenotype, we removed mesoderm of 15 hr post fertilization (hpf) wild type embryos and transplanted, into these hosts, 15 hpf mesoderm grafts expressing GABA_A_Rαβγ or GABA_A_Rα. Only axons that contacted GABA_A_Rαβγ-expressing myocytes in the grafts expressed GABA and ChAT, in addition to SV2 (**Suppl Fig. 14**). Thus, stabilization of GABA depended on these myocytes and not on GABA expression within the spinal cord.

To assess the functional consequence of anatomical innervation by nerve terminals expressing both ChAT and GABA, we recorded mEPPs from GABA_A_Rαβγ-expressing myocytes at 4 dpf. We observed two classes of mEPPs, distinguished on the basis of their rise and decay times and their frequency^5^, with a mean overall frequency of 0.8±0.1 sec^-1^. Those with faster times occurred at a mean frequency of 0.6±0.0 sec^-1^ and were blocked by pancuronium, showing they depended on AChR. Those with slower times occurred at a mean frequency of 0.2±0.0 sec^-1^ and were blocked by bicuculline (**Fig. 3, a** to **f**), indicating they depended on GABA. In contrast, only a single class of mEPPs at a frequency of 1.0±0.1 sec^-1^ was observed when recording from myocytes that expressed GABA_A_Rα.. These mEPPs had rise and decay times similar to mEPPs recorded from GABA_A_Rαβγ-expressing myocytes in the presence of bicuculline (**Fig. 3, g** to **j**). The results of these recordings demonstrate that GABA can be functionally released from motor nerve terminals and innervate myocytes that express GABA_A_Rαβγ receptors.

**Fig 3.**
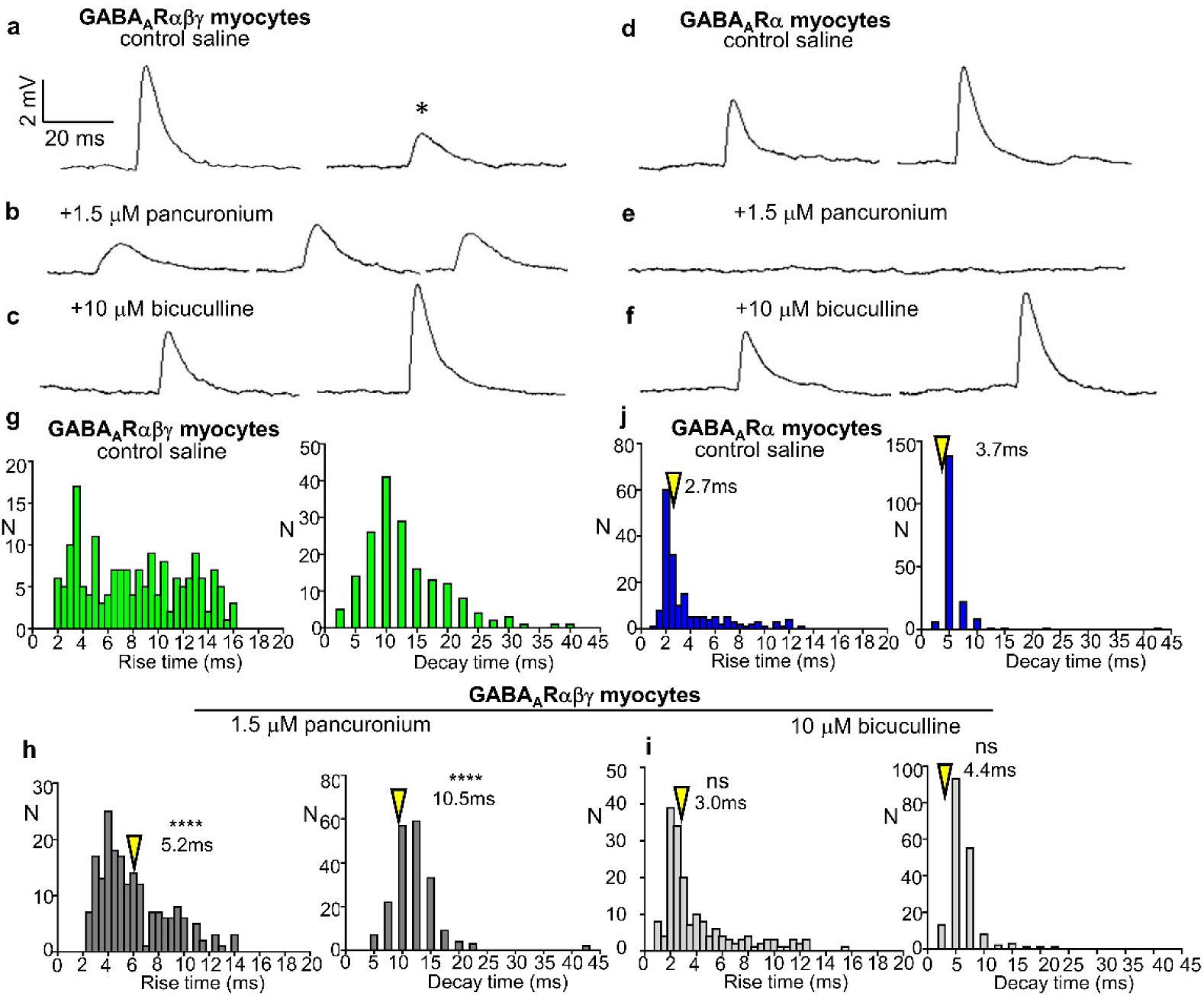
Neuromuscular junctions expressing GABA_A_Rαβγ generate GABAergic and cholinergic mEPPs. (**a-c**) Recordings from GABA_A_Rαβγ myocytes of 4dpf larvae reveal slow-rise and slow-decay GABA_A_R-mediated mEPPs (asterisk) that are pancuronium-resistant and bicuculline-sensitive, as well as pancuronium-sensitive and bicuculline-resistant mEPPS with rise and decay times similar to those described for nicotinic receptor mediated mEPPs. (**d-f**). Rise and decay time distributions for mEPPs in GFP+ myocytes of GABA_A_Rαβγ larvae in presence of saline, pancuronium and bicuculline. (**g-i**) Recordings from GABA_A_Rα myocytes reveal only pancuronium-sensitive mEPPs with kinetics of nicotinic receptor mEPPs. (**j**) Rise and decay time distributions for mEPPs in GFP+ myocytes of GABA_A_Rα larvae in presence of saline. N, number of mEPPs. >178 mEPPs (n≥5 larvae) for each group. Only mEPPs with decay times fit by single exponentials were included. Resting potentials were held near −60 mV. Arrowheads indicate median values. Kolmogorov-Smirnov test comparing rise time and decay time in (**j**) with respective rise and decay time in (**h**) and (**i**). ****p<0.0001, ns not significant.

### Signal transduction by trans-synaptic bridges

We then considered the role of trans-synaptic bridges in retrograde signaling by postsynaptic receptors that regulates the stability of presynaptic transmitters^27–29^. Presynaptic neurexins and postsynaptic neuroligins and dystroglycans are synaptic adhesion molecules (SAMs) that are important for proper maturation and function of synaptic contacts^30^. They bridge the synaptic cleft and create a potential pathway for retrograde signaling. However, in synapse formation assays, SAMs induced excitatory and inhibitory presynaptic specializations at the same time^31^ and did not exhibit the specificity needed if trans-synaptic bridges were to be relevant to our findings. Absent from these assays were the transmitter receptors and their auxiliary subunits and co-receptors. The GABA_A_Rαβγ auxiliary subunit GARLH4 mediates interaction between the γ subunit of the GABA_A_Rαβγ receptor and neuroligin 2, which binds to neurexin^32^. Postsynaptic expression of GABA_A_Rαβ resulted in receptors with functional properties similar to those of GABA_A_Rαβγ^24^, but failed to stabilize presynaptic GABA. We observed that 0 of 12 GABA_A_Rαβ-expressing larvae expressed GABA in motor neuron axons contacting GFP-labeled myocytes and GABA expression remained restricted to the spinal cord (**Suppl Table 2**). This finding indicated that the GABA_A_R γ subunit of the GABA_A_Rαβγ receptor might act through a trans-synaptic bridge to achieve presynaptic stability of the GABAergic phenotype.

To investigate directly whether trans-synaptic bridges stabilize presynaptic transmitters, we first confirmed the presence of neurexin, neuroligin (**Suppl Fig. 15**) and dystroglycans^33^ at the *Xenopus* neuromuscular junction. Knocking down GARLH4 in myocytes, by injecting morpholinos into the V blastomeres (**Suppl Fig. 16** and **Suppl Fig. 17**), prevented the GABA_A_Rαβγ-mediated stabilization of GABA in motor neurons (**Fig. 4, a** to **d**). This result suggested that the GABA_A_R γ subunit of the GABA_A_Rαβγ receptor mediates the expression of presynaptic GABA through GARLH4-neuroligin-neurexin trans-synaptic bridges. Similarly, disrupting AChR-rapsyn-Lrp4-musk-dystroglycan-neurexin trans-synaptic bridges^34,35^, by injecting morpholinos into the V blastomeres to knock down the postsynaptic AChR co-receptor protein, Lrp4^36,37^ (**Suppl Fig. 18**), recapitulated the destabilization and loss of ChAT that we observed in the presence of pancuronium (**Fig. 4, e** to **l**). Larvae in which Lrp4 was knocked down exhibited two classes of mEPPs at 4dpf, differing in their rise and decay times (**Suppl Fig. 19**) and their frequency, with an overall mean mEPP frequency of 0.7±0.1 sec^-1^. Pancuronium-resistant NBQX-sensitive glutamatergic mEPPs were had a mean frequency of 0.5±0.0 sec^-1^. Pancuronium-sensitive, NBQX-resistant cholinergic mEPPs had a mean frequency of 0.1±0.0 sec^-1^. These cholinergic mEPPs, for which there were also fewer AChR^38^, were smaller in amplitude compared to cholinergic mEPPs in larvae expressing the control morpholino (1.7±0.1 mV vs 2.7±0.1 mV; n^mEPP^=177, N^larvae^=7; p<0.0001, two-tailed t-test). mEPPs in control larvae implanted with a saline bead occurred at a mean frequency of 1.2±0.1 sec^-1^. Pancuronium may phenocopy the effect of knocking down Lrp4 by binding to the α and γ subunits of the AChR^37^ and altering the AChR-Lrp4 interaction, thereby disrupting signaling through cholinergic trans-synaptic bridges.

**Fig 4.**
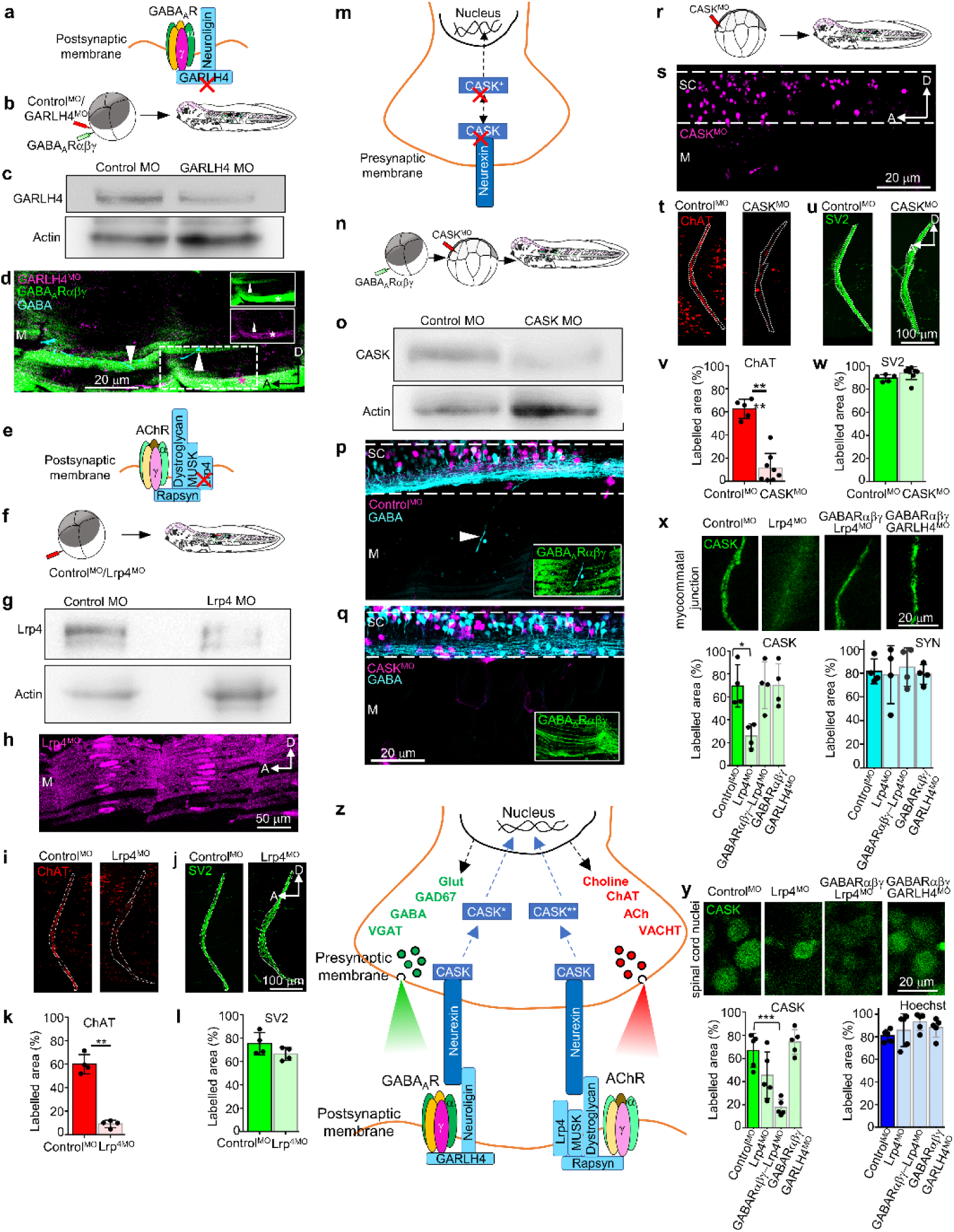
Receptor-driven neurotransmitter stabilization is mediated by trans-synaptic bridges. (**a**) GARLH4 links GABA_A_Rαβγ to neuroligin in the postsynaptic membrane. (**b**) Staggered injection of ventral blastomeres (V) with GABA_A_Rαβγ mRNA followed 1 minute later by low dose of control^MO^ or GARLH4^MO^ (3 nl of 1 mM MO). (**c**) Presence of GARLH4 and morpholino knockdown validation by Western blot. (**d**) Axons contacting GABARαβγ myocytes express GABA (arrowheads; 5/5). Axons contacting a GARLH4^MO^ GABA_A_Rαβγ myocyte do not express GABA (asterisk, 0/12). *Insets*, individual channels for GFP (GABARαβγ) and GARLH4_MO_. Dashed box shows the area depicted in the inserts. 3dpf. (**e**) Lrp4 links to AChR through rapsyn and to MUSK and dystroglycan in the postsynaptic membrane. (**f**) Injection of ventral blastomeres (V) with a high dose of Lrp4_MO_ or control_MO_ (6 nl of 1 mM MO). (**g**) Morpholino knockdown validation by Western blot. (**h**) Lrp4_MO_ expression in the myotome. (**i, j**) Myocommatal junctions of larvae with control or Lrp4_MO_ stained for ChAT and SV2. (**k, l**) Intensity of expression of ChAT and SV2. n=4, **p=0.0092, two-tailed *t-test*. 3dpf. (**m**) CASK links to neurexin. (**n**) Injection of GABARαβγ RNA in ventral blastomeres (V) and a high dose of CASK_MO_ (6 nl of 1 mM MO) in dorsal blastomeres 1.2 (D1.2). (**o**) Presence of CASK and morpholino knockdown validation by Western blot. (**p**) GABA+ axon in myotome of control_MO_ GABARαβγ larva (arrowhead, 6/6). (**q**) 3dpf CASK^MO^ GABA_A_Rαβγ larvae do not show GABA expression in the myotome (0/12). (**r**) Injection of CASK^MO^ in dorsal blastomeres 1.2. (**s**) larva expressing CASK_MO_ in the SC. (**t, u**) Myocommatal junctions of larvae with control_MO_ or CASK_MO_ stained for ChAT and SV2. (**v, w**) Intensity of expression of ChAT and SV2. n≥5, ****p<0.0001, two-tailed *t-test*. (**x, y**) Myocommatal junctions of larvae with control or Lrp4^MO^ or GABA_A_Rαβγ-Lrp4^MO^ or GABA_A_Rαβγ-GARLH4^MO^ stained for CASK in synaptophysin (SYN)+ neuronal processes along the myocommatal junction and in Hoechst+ nuclei in the spinal cord. (**x**) Intensity of expression of CASK and SYN. N=4, *p<0.01, one-way ANOVA (F_3,12_=6.316, p=0.0081). (**y**) Intensity of expression of CASK and Hoechst. N=5, ***p<0.0001, one-way ANOVA (F_3,12_=16.5, p<0.0001). 3dpf. (**z**) Summary of interactions between neurotransmitter receptors and transsynaptic bridge proteins that convey retrograde signals to achieve presynaptic transmitter stabilization. n/N, larvae with GABA+ axon/total larvae observed. Red X in **a, e** and **m**, protein knocked-down. M, myotome; SC, spinal cord; NT, neurotransmitter; D, dorsal; A, anterior.

We next considered CASK (Ca^2+^/calmodulin-activated Ser-Thr kinase) as a candidate that could receive signals from the presynaptic ends of the trans-synaptic bridges and stabilize transmitter expression in the motor neurons. CASK is a presynaptic membrane-associated guanylate kinase^39^ and transcription factor that binds to neurexin protein (**Fig. 4m**) and is present at both glutamatergic^40^ and GABAergic synapses^41^. CASK pre-mRNA is subject to alternative splicing that yields proteins with preferences to interact with many targets^42^. Autophosphorylated CASK translocates to the nucleus and induces transcription of genes essential for development^43,44^. Additionally, neurexin-1 competes as a CASK phosphorylation substrate, preventing CASK autophosphorylation^45^. The splice variants, together with differential phosphorylation, identified CASK as a potentially significant player in stabilizing presynaptic expression of different neurotransmitters. We found that CASK is expressed in the myotome and spinal cord of wild type *Xenopus* larvae (**Fig. 4, x** and **y;** and **Suppl Fig. 20**). Knocking down presynaptic CASK, by injecting morpholinos into the D1.2 blastomeres to target the spinal cord (**Suppl Fig. 20**), disrupted both AChR-mediated ChAT stabilization and GABA_A_Rαβγ-mediated GABA stabilization in motor neurons (**Fig. 4, n** to **w;** and **Suppl Table 1**). Knockdown of CASK, Lrp4 or GARLH4 did not alter laminin expression or the extent of labelling by synaptophysin (SYN), indicating there was no change in the gross morphology of the myocommatal junction or postsynaptic myocytes (**Fig. 4, u** and **w** and **x;** and **Suppl Fig. 21**). However, presynaptic expression of CASK in spinal nuclei and in myocommatal junctions was dependent on transmitter receptor identity **(Fig. 4, x** and **y**). In control larvae, CASK expression was distributed between the nuclei and the myocommatal junctions. Disrupting the cholinergic trans-synaptic bridge by knocking down Lrp4 reduced CASK expression to 30% of controls in the SYN-labeled myocommatal junctions, but not in the nuclei. Introducing the GABA_A_Rαβγ−mediated GABAergic trans-synaptic bridge and simultaneously disrupting the cholinergic trans-synaptic bridge reduced CASK expression in the Hoechst-labeled nuclei to 20% of controls, but not in the SYN-labelled myocommatal junctions. Disrupting the GABAergic trans-synaptic bridge in the presence of the cholinergic trans-synaptic bridge recapitulated the CASK expression observed in control larvae.

## Discussion

Our results demonstrate that postsynaptic neurotransmitter receptors regulate the stability of presynaptic neurotransmitters at newly formed neuromuscular junctions. This regulation is achieved by retrograde signaling that operates through trans-synaptic bridges (**Fig. 4**). When one bridge is disrupted by knock down of Lrp4, transmitter synthesis is reduced. When another bridge is introduced by expression of GABA_A_Rαβγ, transmitter synthesis is stabilized. Neurexin serves as a presynaptic link of the trans-synaptic bridges, receiving postsynaptic receptor-dependent signals and transmitting them through the neurexin-interacting protein CASK to achieve presynaptic cholinergic or GABAergic stabilization. The receptor-dependent distribution of CASK within the presynaptic neuron, coupled with its roles in scaffolding the synapse^46^, organizing presynaptic voltage-gated calcium channels^47,48^ and regulating neuronal gene transcription^43^, provides a mechanism by which different postsynaptic receptors stabilize cognate neurotransmitter expression through trans-synaptic bridges. This signaling system of trans-synaptic bridges shares features with clustered protocadherin cell adhesion molecules^49^, some of which change the nuclear versus cytoplasmic distribution of β-catenin to regulate the Wnt pathway^50^.

The reduction in presynaptic transmitter markers upon blockade of postsynaptic receptors is consonant with features of presynaptic homeostatic plasticity, in which a compensatory increase in neurotransmitter upon loss of receptor function is preceded by a compensatory change in receptor subunits^14,15^. In *Xenopus* myocytes, where the AChR block cannot not be rescued by alternative subunits^16^, presynaptic synthesis of ACh was reduced and expression of glutamate receptors in myocytes^5,51^ led to stabilization of presynaptic glutamate^52,53^.

The demonstration that trans-synaptic bridges are involved in transmitter stabilization does not preclude a role for postsynaptic diffusible factors, some of which have been shown to influence synapse formation and maintenance retrogradely^54–57^. Future work will determine whether deficits in postsynaptic receptors or trans-synaptic bridges contribute to reduced transmitter levels at neuronal synapses in the mature nervous system and to pathological change in neurological disorders.

## METHODS

### Animals

Frogs were purchased from Nasco and Xenopus 1 and maintained on the daily light-dark cycle. Experiments were performed throughout the year over a period of 4 years. No seasonal differences in the data were observed. Adult female frogs were primed with 10 IU human chorionic gonadotropin (HCG; Millipore Sigma) 1 week prior to collection of eggs. Testes were harvested and stored in 20% fetal bovine serum or were cryopreserved till fertilization^58^. Ovulation was induced by injection of 500 IU HCG 12 hr prior to egg collection and fertilization. Eggs were fertilized *in vitro* and embryos developed in 0.1X Marc’s modified Ringer’s solution (MMR: 100 mM NaCl, 2 mM KCl, 1 mM MgSO_4_, 5 mM HEPES, 0.1 mM EDTA, 2 mM CaCl_2_, pH 7.8). Developmental stages were determined from developmental tables^59,60^. All animal procedures were performed in accordance with institutional guidelines and approved by the UCSD Institutional Animal Care and Use Committee.

### Local drug delivery

Spatial and temporal control of delivery of pharmacological agents was achieved using agarose beads (100-200 mesh, Bio-Rad)^8,21^. Beads were washed in 2 mM Ca^2+^ medium for 3 h at 22°C, replacing washing solution every 20 min, and loaded for 2 h at 4°C on a bi-directional rotator with 2 mM Ca^2+^ medium containing drugs 30 μM pancuronium (Fisher Scientific), or 38.9 μM curare (Sigma-Aldrich), or saline (0.9% sodium chloride; stock solutions of drugs were diluted in saline). Drug concentrations were 10X the concentration tested in myocyte cultures^4,5^, to compensate for dilution during diffusion from the bead. Beads of 120 μm diameter were selected and implanted into the mesoderm (120 µm from the neural tube) ∼400 μm behind the eye primordium at 20 hr post fertilization (hpf) (Stage 18), allowed to develop in 0.1X MMR to 2 days post fertilization (dpf) (Stage 35-36), 3 dpf (St 41), 4 dpf (St 45) or 7 dpf (St 48), and processed for immunohistochemistry. Each experimental group contained 4-8 bead-implanted embryos. Only myocommatal junctions ≤200 μm from a bead were considered for analysis.

### Plasmids and morpholinos

Expression of GABA_A_R was achieved with GABA_A_R subunit plasmids for α1-EGFP, β2 and γ2 generated in the Czajkowski lab^26^. The GABA_A_R subunits were subcloned into pCS2 by VectorBuilder (Santa Clara, CA)-pCS2-GABAr alpha-GFP VB180118-1043eef, pCS2 GABAr beta VB180303-1004bat, pCS2 GABAr gamma VB180303-1005aab. The lissamine-tagged GARLH4, Lrp4 and CASK morpholinos were supplied by GeneTools (Philomath, OR) (GARLH4 MO1 5’-TTCTTGTGAAGTCAACATCTTGG-3’, GARLH4 MO2 5’-GAGTCCTCACTCAGTATCC-3’, GARLH4 negative control MO 5’-GGTTCTACAACTGAAGTGTTCTT-3’, Lrp4 MO 5’-CCCATAATGCCACTTCTCC-3’, Lrp4 negative control MO 5’-CCTCTTCACCGTAATACCC-3’, CASK MO 5’-GGGTTTAACACCACCACAAAGGA-3’, standard control oligo 5’-CCTCTTACCTCAGTTACAATTTATA-3’).

### Microinjections

Plasmids were amplified with the QIAprep Spin miniprep kit (Qiagen) and linearized with Not1-HF (New England Biolabs). α1-EGFP, β2 and γ2 capped mRNA were transcribed using the mMessage mMachine SP6 kit (Invitrogen). The integrity of the mRNA was assessed using both Nanodrop and agarose gel electrophoresis. On the day of microinjections, eggs were fertilized in 0.1X MMR and 20 min later treated in 2% cysteine (Sigma-Aldrich) to remove the jelly coat. They were rinsed 10 times in 0.1X MMR and allowed to develop to the 4-cell stage in 0.1X MMR at 25°C. Embryos were transferred to 6% Ficoll (Sigma-Aldrich) for injections. Using a Picospritzer III, α1, β2 and γ2 mRNA were co-injected into both ventral blastomeres (V) in 1:1:1 ratio along with phenol red (to visualize the solution), to a total volume of 3 nl per blastomere, to target expression to myocytes^61^. 6 nl of 1 mM Lrp4 MO and 1 nl or 3 nl of 1 mM GARLH4 MO or their respective control morpholinos were injected into the ventral blastomeres to knockdown the respective mRNAs. 6 nl of 1 mM CASK MO or its control morpholino were injected into dorsal blastomeres 1.2 (D1.2) at the 16-cell stage to target expression in the spinal cord^25^. Higher doses of Lrp4 MO and CASK MO were delivered to achieve widespread knock-down of the Lrp4 and CASK, as strong changes in phenotype were not observed at 3 nl dose. 3 nl of 1mM GARLH4 labelled almost all myocytes. This dose along with GABARαβγ mRNA injection was used to test the necessity of GARLH4, while the low dose of 1 nl GARLH4 MO followed 1 minute later by GABARαβγ mRNA injection was used to achieve sparse labelling of myocytes with the morpholino. Injected embryos were transferred to 0.1X MMR and developed at 25°C. Lissamine was detected immediately post-injection, while GFP fluorescence was detected at 17 hpf (St 15) as previously reported^62^, and persisted up to 7 dpf (St 48).

### Whole mount immunohistochemistry

Whole *Xenopus* larvae were fixed at the indicated stages of development in 4% PFA and 0.1% glutaraldehyde for 1 h. After 3 washes in 1X PBS, the epidermis and head of fixed larvae were removed in 1X PBS and the larvae were transferred into a 12-well plate containing blocking solution (5% normal donkey serum, 1% fish gelatin, 0.5% Triton-100 in 1X PBS) for 1 h. Larvae were then transferred to fresh wells containing primary antibodies diluted in 50% blocking solution in 1X PBS and incubated at 4°C for 3 d. Antibodies used were ChAT (1:500, Millipore, AB144P), SV2 (1:500, Developmental Studies Hybridoma Bank, AB2315387), VGLUT1 (1:200, Millipore, AB5905), GLUR1 (1:200, MilliporeSigma, MAB2263), NR1 (1:200, MilliporeSigma, MAB363), GABA (1:200, Millipore, ABN131), VGAT (1:300, cytoplasmic domain, Synaptic Systems, 131013), GFP (1:500, Synaptic Systems, 132005), Hb9 (1:10, Developmental Studies Hybridoma Bank, 81.5C10), GAD67 (1:250, Millipore, MAB5406), Glycine (1:200, Millipore, AB139), Lim3 (1:200, Millipore, AB3202), rAldh1a2 (1:8000, ZMBBI Columbia, CU1022), pan-Neurexin (1:200, kind gift from Peter Scheiffele), pan-Neuroligin (1:200, Invitrogen, PA5-77523), synaptophysin (1:200, Synaptic Systems, 101002), laminin (1:200, MilliporeSigma, L9393), Lrp4 (1:500, extracellular domain, generous gift from Stephan Kröger), GARLH4 (1:200, LifeSpan Biosciences, LS-C170010) and CASK (1:200, Santa Cruz Biotechnology, SC-13158). After 3 washes in 1X PBS, larvae were incubated in the dark at 4°C for 24 h with secondary antibodies (1:500, Alexa Fluors 488, 555, 594 and 647, Jackson Immunoresearch). For CASK immunostaining experiments larvae were incubated in Hoechst stain (1:8000, MilliporeSigma, B2883)) for 30 mins. Following 3 washes with 1X PBS, larvae were serially dehydrated in 50% methanol and 100% methanol for clearing with BABB (Benzyl alcohol:benzyl benzoate 1:2, Sigma-Aldrich) for 24 h at 4°C. After 24 h the larvae were transferred to wells containing fresh BABB and reincubated for 24 h. Larvae were mounted onto slides with the side exhibiting maximum expression facing up. Confocal images of whole larvae were acquired on a Leica Stellaris 8 confocal microscope with 25x/0.95 water immersion objective, at a z-resolution of 0.5 μm, and analyzed with FIJI. For the CASK-Hoechst immunostaining experiments, images of whole larvae were acquired on the Leica SP8 with 40X/1.30 oil immersion objective, at a z-resolution of 0.5 µm and analyzed with FIJI. Imaging parameters were kept constant within each experiment.

### Image analysis

Analysis of axons running down myocommatal junctions: For bead-implantation experiments, the myocommatal junction ≤200 µm anterior or posterior to the bead was analyzed. For each larva the first optical slice exhibiting SV2 expression at the myocommatal junction was determined and optical slices were stacked from that slice onwards to obtain 25 μm thick maximum intensity projections from 50 slices, each of 0.5 μm thickness. These projections enabled examination without contamination of the SV2 signal along myocommatal junctions by SV2 processes in the spinal cord. The regions of interest (ROIs) for the myocommatal junctions were outlined with the FIJI freehand selection tool using the SV2 signal as reference. Each maximum intensity projection image was filtered for 2x background SV2, ChAT or VGLUT1 mean signal intensity and this threshold was applied to all slices of the stacks. The SV2 and ChAT or VGLUT1 images were converted into 8-bit format and then into binary images. The percentage area in the ROI labelled with the signal was measured for SV2 and ChAT or VGLUT1. Data analysis parameters were kept constant within each experiment, allowing aggregation of data from independent batches of larvae. For CASK experiments the ROIs for the myocommatal junctions were outlined with the FIJI freehand selection tool using the synaptophysin signal as reference, while the ROIs for nuclei were outlined with the FIJI freehand selection tool using the Hoechst signal as reference.

Analysis of glutamate receptors on myocytes between two myocommatal junctions, spanning one chevron: For each larva the first optical slice exhibiting GLUR1 or NR1 staining was determined, and optical slices were stacked from that slice onwards to generate 25 μm thick maximum intensity projections; these enabled examination of the myocytes in a chevron without the appearance of spinal cord that could otherwise obscure visibility of the axons of motor neurons. The percentage labelled area analysis of whole mount immunostained preparations included myocytes spanning one chevron immediately caudal to the bead, but excluded tissue edges, damaged cells, residual skin and pigment cells. Data were analyzed as described above.

Analysis of axons expressing GABA along the A-P axis: For GABA receptor expression experiments, each larva was imaged at 4 positions along the A-P axis to cover its entire length (head to tail). At each position, ∼200 0.5 μm optical slices (not maximum intensity projections) were acquired, thus covering the entire dorsal-ventral axis. These slices were examined one-by-one to determine the number of GABA-expressing axons in the myotome. Twenty 3 dpf GABARαβγ-expressing larvae were examined in this manner. When a terminal axon expressing GABA was detected in the myotome of a larva, its chevron position was determined, and the position of the axon was marked on a schematic of a 3 dpf larva using the ImageJ multipoint tool. These points were saved as a ROI. All the ROIs were overlaid together on the 3 dpf schematic to determine if there was an A-P distribution preference for these axons in the myotome. Chevrons along the schematic were broadly divided into three groups for the purpose of analysis^63^: anterior (light blue), chevrons 1 to 7; middle (blue), chevrons 8 to 16; and posterior (dark blue), chevrons 17 to 21. Chevron 21 is a composite of chevrons including and following chevron 21 in the tail, which are harder to discern.

Analysis of GABA+Hb9+ cell bodies in the spinal cord: Images of the larvae were acquired as described above. A 3D-image was created from the 0.5 μm slices. The GABA+ axon was located in the myotome and traced back to the spinal cord using the 3D-Viewer and Simple Neurite Tracing plugins in ImageJ.

Analysis of GABA, VGAT and SV2 colocalization: Colocalization was determined for each larva by examining all the optical sections within a confocal stack without maximal projection. Processes expressing the markers-GABA, VGAT and SV2 in at least 3 consecutive z-planes were termed colocalized. The entire confocal stack was scanned optical section-by-optical section for myocytes of interest, i.e. a GFP+ myocyte along the same D-V axis as the colocalized processes.

In each larva the first optical slice exhibiting GFP expression in myocyte of interest was determined and optical slices were stacked from that slice onwards to obtain 10 μm-thick maximum intensity projections from 20 slices, each of 0.5 μm thickness.

### Cell counting

FIJI software was used to count fluorescent, immunolabeled ChAT+, Hb9+, Lim3+, GABA+, and GAD67+ cell bodies in the spinal cord; all slices within the confocal stack through the spinal cord were examined without maximal projection. Representative images are maximum intensity projections of 5 consecutive slices.

### Protein extraction and ELISA

ELISA assays for GFP-tagged GABA_A_Rα subunit expression in the muscle and spinal cord were performed per the manufacturer’s instructions (GFP ELISA Kit, Cell Biolabs). Larvae were anaesthetized in 0.02% tricaine methanesulphonate (10 g/L stock solution) for 1 min, skinned and gut removed using jewellers’ forceps. Larvae were secured to Sylgard in a 60 mm plastic Petri dish with a minutien pin (dorsal side up). The melanocyte layer covering the dorsal side (spinal cord) was gently removed. 0.05% collagenase was added to the dorsal side of the larva using a glass capillary, for 5 min. The collagenase was then removed and replaced with 1X PBS. The brain and spinal cord were lifted out, collected into a 2 ml Eppendorf and frozen immediately in liquid nitrogen. The trunk myotome was collected in another Eppendorf and frozen the same way. Thirty brains and spinal cords were pooled for a single experiment to yield sufficient protein. Samples were sonicated in RIPA lysis buffer (150 mM NaCl, 50 mM Tris (pH 7.4), 2 mM EDTA, 1% NP-40, 0.1% SDS, 0.1 % (v/v) Protease Inhibitor) and cell debris was pelleted by centrifugation at 5000xg for 10 min. Protein concentration of the supernatant was estimated using the BCA method (ThermoFisher).

### Mesoderm grafting

Myocyte-dependent stabilization of GABA in innervating motor neuron axons was assessed by mesoderm grafts. To obtain myocyte-specific GABA_A_Rαβγ or GABA_A_Rα expression, presomite mesoderm of wild-type embryos 15 hpf (St 13-14) was replaced with presomite mesoderm explant dissected from sibling embryos expressing either GABA_A_Rαβγ or GABA_A_Rα transcripts. The grafting technique was similar to that previously described^64^. Briefly, embryos were manually demembranated with jeweller’s forceps, then incised using a tungsten needle in the posterior region next to the blastopore, at the level of the presumptive tail-forming region. The neural plate was cut laterally, left attached at the midline and lifted using a hair loop to free access to the underlying presomitic mesoderm. The explant of presomitic mesoderm from the host was replaced by a similarly-sized one from a GFP+ embryo, using jeweller’s forceps. To immobilise the graft and promote healing, the operated region was covered by a small glass coverslip bridge. The grafted embryos were allowed to heal under these bridges until they were sealed and then moved to 0.1X MMR for further development.

### Electrophysiology of mEPPs

Sample preparation: 4 dpf (St 43) larvae were anaesthetized in ice cold 10% MMR. After ensuring non-responsiveness to touch, a larva was transferred into a 60 mm Petri dish with ice cold external solution, decapitated and the epidermis peeled back using sharp forceps to expose the trunk myotome. The larva was moved to a Sylgard platform (∼1 mm thick) in the same Petri dish and glued laterally in place with Vetbond tissue adhesive (3M). A glass capillary was used to apply a drop of Vetbond glue to Sylgard. The Sylgard platform with the larva was transferred into a recording chamber containing external solution (NaCl 115 mM, KCl 4 mM, CaCl_2_ 3 mM, MgCl_2_ 1 mM, HEPES 5 mM, D-Glucose 10 mM; pH 7.2, Osm 250-255).

Sharp electrode recording: G150F-3 (borosilicate: standard wall with filament, Warner Instruments) capillaries were loaded with internal solution (3M KCl) through microfil/microloader tips. Filled-capillaries were placed sharp side down on plasticine stuck to the edge of a shelf for gravity-assisted removal of air bubbles. Glass pipettes with resistance of 54-75 MΩ were used for recordings. GFP-labeled green myocytes, identified by the presence of sarcomeres visualized with an upright Zeiss microscope and a water immersion 40X/0.8 objective, were impaled within 40 μm of the myocommatal junction for recordings. Multiclamp 7008, Clampex 10.7 software and Clampfit 10.7 data interfaces were used for acquisition and analysis of miniature postsynaptic end-plate potentials (mEPPs). The membrane potential was held at −60 mV with bridge balance for the duration of recordings. mEPPs were recorded for a duration of 5 min, after which blockers (pancuronium 1.5 μM, bicuculline 10 μM, or NBQX 20 μM) were perfused into the recording chamber (2 ml/min). Following 10 min of perfusion, mEPPs were recorded in the presence of the blockers to determine receptor-dependence. Blockers were judged effective when they reversibly reduced the frequency and amplitude of mEPPs. The drugs were washed out for 20 min and recovery mEPPs recorded to ensure the sample preparation was healthy throughout the recording. The recordings from bead-implanted larvae and morpholino-injected larvae were performed in the presence of 1 μM tetrodotoxin to prevent contribution of off-target effects of pancuronium or NBQX. Rise times were scored from 10% to 90% of peak and decay times were determined by fitting exponentials with the MiniAnalysis program (Synaptosoft Inc, Decatur, GA). The rise and decay time distributions in different conditions were compared using Kolmogorov-Smirnov test and considered significantly different when *P* < 0.05. Frequencies were scored as mEPPs min^-1^.

### Western blotting

Protein for Western blotting of components of trans-synaptic bridges was extracted as described above except that larvae were anaesthetized on ice and RIPA lysis buffer was supplemented with phosphatase inhibitors (PhosSTOP, Roche). 30 μg of total protein were loaded per well and Ponceau S staining was used to check total protein levels on the membrane. Protein detection by Western blotting was performed for Lrp4 (1:8000, extracellular domain, generous gift from Stephan Kröger), GARLH4 (1:500, LifeSpan Biosciences, LS-C170010) and CASK (1:500, Santa Cruz Biotechnology, SC-13158). The same blots were re-probed for the loading control actin (1:1000, Sigma-Aldrich, A2066-100UL). Signal was detected using Clarity ECL Western substrate (BioRad) and imaged on a BioRad Chemidoc Touch Imaging System.

### Statistics

Values were expressed as mean ± SD. Statistical differences were analyzed using the unpaired, two-tailed Student’s *t-test* or one-way ANOVA or Kolmogorov-Smirnov test. The statistical test used and the P values for each measurement are provided in the figure legends. *P*<0.05 was considered statistically significant. All statistical analysis was performed in GraphPad Prism.

## Data availability statement

The data that support the findings of this study are available from the corresponding author upon request.

## Acknowledgements

We thank all members of the Spitzer laboratory for discussions and critical feedback; K. Marek for discussions of acknowledgement signals; I. Gregor and R. Aricescu for discussions of receptor pharmacology and trans-synaptic bridges; C. Kintner for advice on *Xenopus* blastomere lineage; A. Ray and E. Park for guidance on miniature analysis; A. Glavis-Bloom, S.U. Choi and S. Malladi for technical assistance; D.K. Berg and L.R. Squire for comments on the manuscript. This work was supported by NSF 2051555 and the Overland Foundation. Some of the microscopy utilized the UCSD School of Medicine Microscopy Core, supported by NIH grant NS047101.

## Author contributions

S.K.G and N.C.S conceived the study and designed the experiments with consultation and advice from H-q.L. and M.P. S.K.G. performed all the experiments. M.H. and H.T.C. performed electrophysiology experiments with S.K.G. Y.I. and L.S. contributed to the identification of motor neuron identity. J.B. and C.C. contributed GABA_A_ receptor clones and advice regarding expression. J.B.L and L.N.B provided instruction in embryonic tissue transplantation. S.K.G. and N.C.S wrote the manuscript with contributions from all authors.

## Competing interest statement

The authors declare no competing interests.

## Additional information

1. Supplementary information is available for this paper.
2. Correspondence and requests for materials should be addressed to Swetha K. or Nicholas C.
3. Peer review information Completed by Nature staff
4. Reprints and permission information are available at www.nature.com/reprints

## Notes

### Competing Interest Statement

The authors have declared no competing interest.

### Summary of Updates

1) Figures 1,3 and 4 revised. 2) Section on "Blocking endogenous acetylcholine receptors at the neuromuscular junction" and Discussion section updated to place the findings in context of previous studies on presynaptic homeostatic plasticity. 3) Updated reference list. 4) Supplemental files updated

