## Supplementary Materials for "Postsynaptic receptors regulate presynaptic transmitter stability through trans-synaptic bridges"

### Supplementary Material

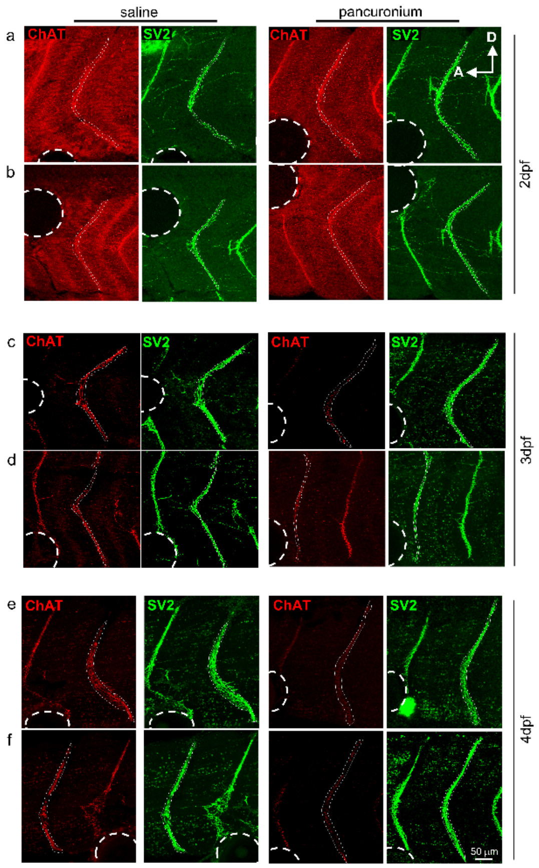

**Suppl Fig 1. Local pancuronium block of AChR in myocytes reduces ChAT expression in innervating axons.** ChAT and SV2 staining of larvae implanted with saline- or pancuronium-loaded beads at 19 hpf and examined at 2dpf (**a-b**), 3dpf (**c-d**) and 4dpf (**e-f**). Dashed circles indicate positions of agarose beads. Dotted lines outline myocommatal junctions analyzed. A, anterior; D, dorsal.

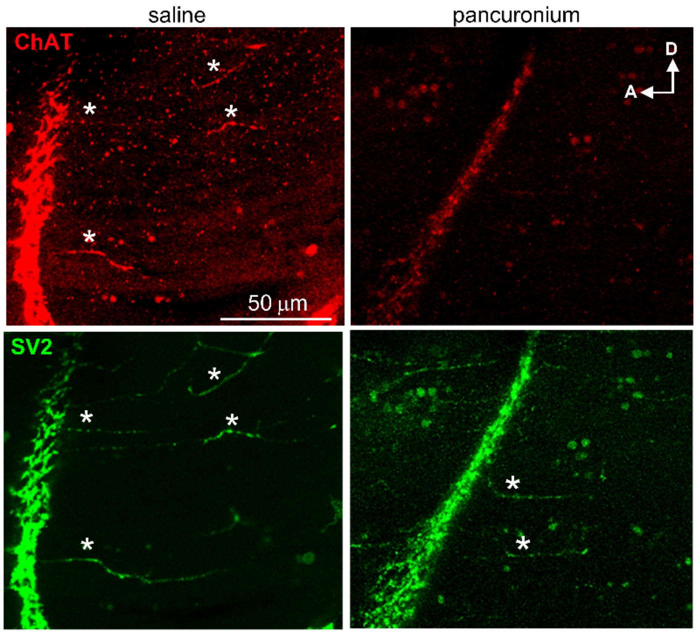

**Suppl Fig 2. Local pancuronium block of AChR reduces ChAT expression in nerve terminals on the myotome.** SV2+ processes seen at the myocommatal junction and in the myotome (asterisks) are ChAT+ in saline but not in pancuronium-loaded agarose bead-implanted larvae at 4dpf. A, anterior; D, dorsal.

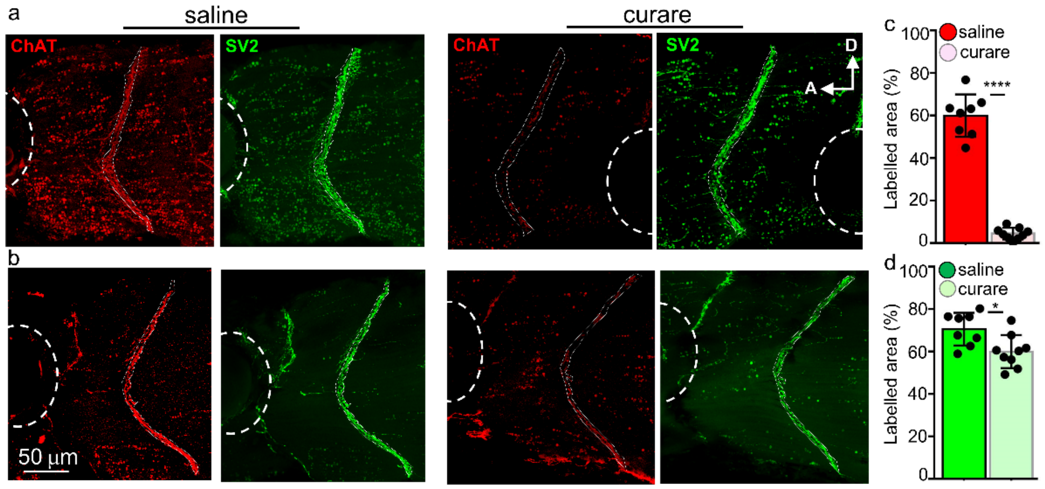

**Suppl Fig 3. Local curare block of AChR recapitulates reduction in ChAT expression in innervating axons seen in the presence of pancuronium.** (**a-b**) Two examples of ChAT and SV2 staining at 4dpf in larvae implanted with saline- or curare-loaded agarose beads at 19hpf. Dashed circles indicate positions of beads. Dotted lines outline myocommatal junctions quantified in (**c**) and (**d**) for ChAT and SV2 respectively. ****$p<0.0001$ and *$p=0.0130$ with unpaired t-test. $n\geq8$ myocommatal junctions from $\geq8$ larvae per group. A, anterior; D, dorsal.

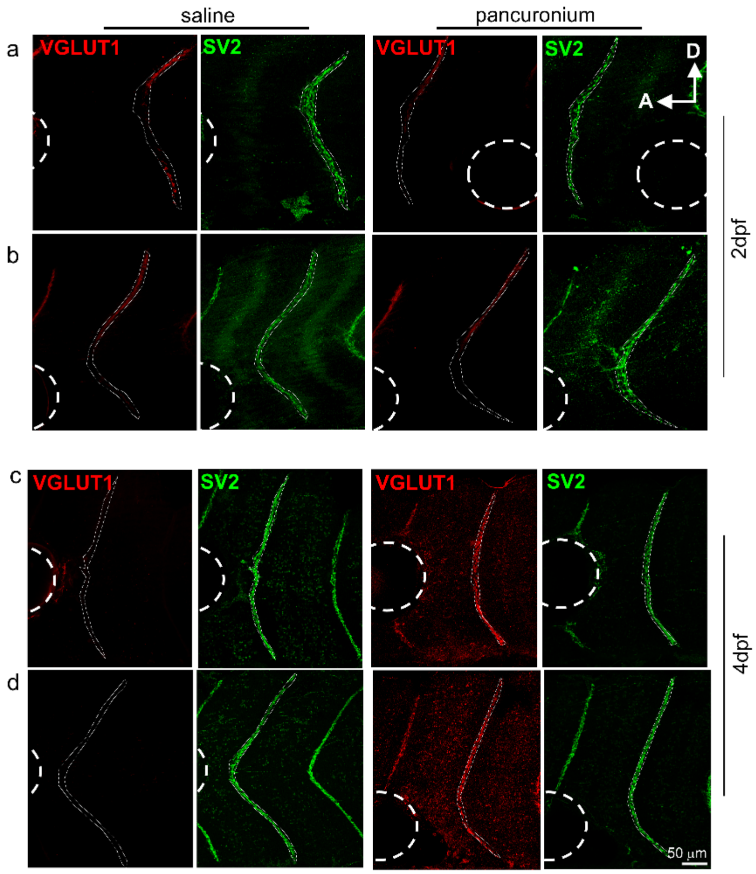

**Suppl Fig 4. Local pancuronium block of AChR in myocytes increases VGLUT1 expression in innervating axons.** Two larvae implanted with saline- or pancuronium-loaded agarose beads at 19hpf and examined at 2dpf (**a-b**) and 4dpf (**c-d**) for VGLUT1 and SV2. Dashed circles indicate positions of agarose beads. Dotted lines outline myocommatal junctions. A, anterior; D, dorsal.

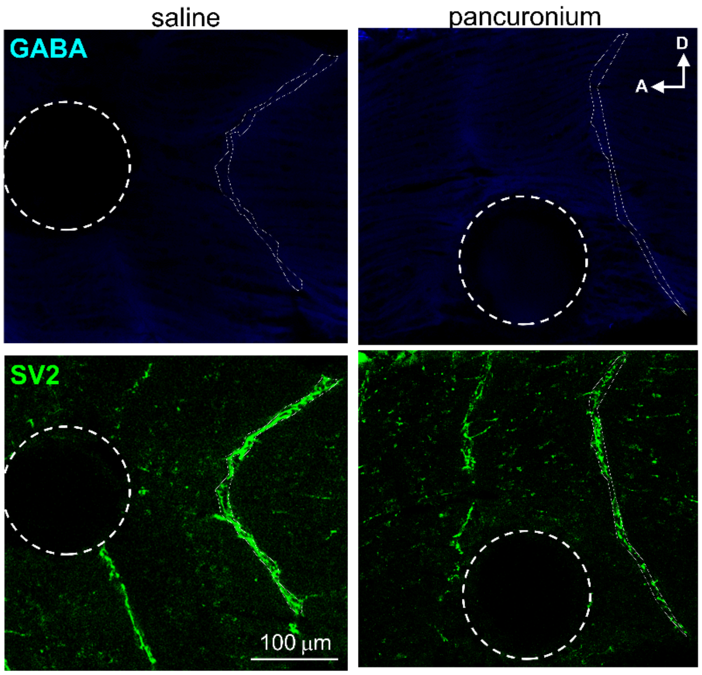

**Suppl Fig 5. Local pancuronium block of AChR does not lead to GABA expression in innervating axons.** GABA is not detected in the myotome of saline- and pancuronium-loaded agarose bead-implanted larvae at 3dpf. White dashed circles indicate positions of agarose beads. A, anterior; D, dorsal.

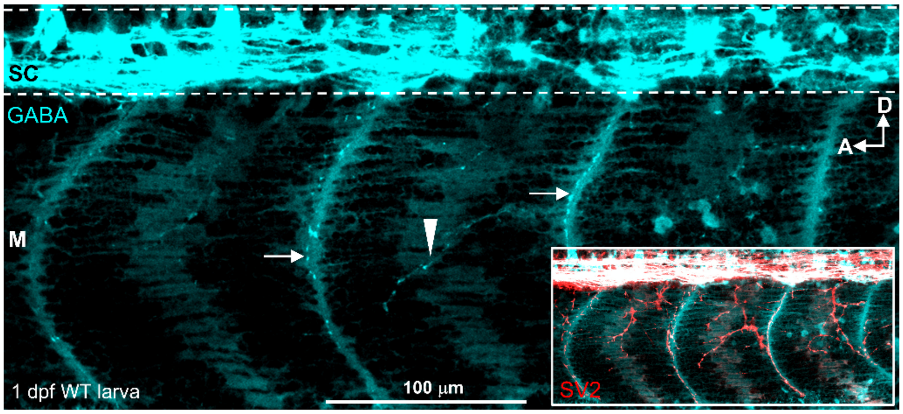

**Suppl Fig 6. GABA expression in axon terminals in myocommatal junctions and on myocytes early in embryonic development.** 1 dpf GABA expression is detected along myocommatal junctions (arrows) and in axons innervating the myotome (M, arrowhead) of a wild type larva. Inset shows GABA and SV2 expression. A, anterior; D, dorsal. SC, spinal cord.

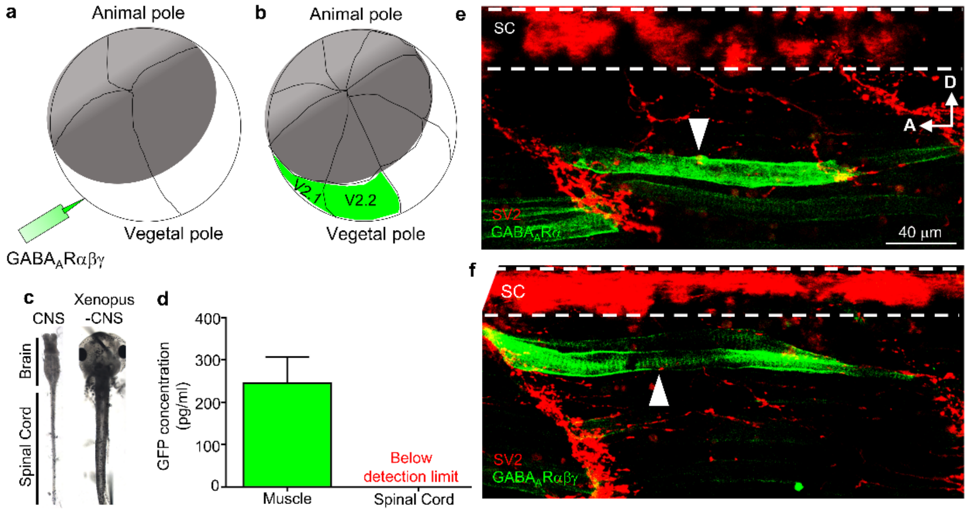

**Suppl Fig 7. Targeting the muscle-specific ventral blastomeres for $GABA_A$ receptor misexpression in myocytes.** (**a**) Schematic of the *Xenopus* embryo injected at the 4-cell stage with $GABA_AR\alpha\beta\gamma$ in the ventral blastomeres (V) towards the vegetal pole. (**b**) Schematic of *Xenopus* embryo at the 16-cell stage with $GABA_AR\alpha\beta\gamma$ targeted only to blastomeres V2.1 and V2.2 (green) that contribute to somites but not to the nervous system. (**c**) Images of the dissected CNS (brain + spinal cord) of a 3dpf larva. Spinal cords (SC) from these dissections were processed for ELISA. (**d**) ELISA for GFP-tagged $GABA_AR\alpha$ subunit expression in the muscle and SC after injection of the V blastomere with GFP-$GABA_AR\alpha$ mRNA shows GFP protein in the muscle sample but not in the SC sample (detection threshold 30 pg/ml). **(e, f)** Examples of sparse $GABA_AR\alpha$-only expression (**e**) and $GABA_AR\alpha\beta\gamma$ expression (**f**) in myocytes (white arrowheads) but not in the spinal cord (SC). SV2 labels axon terminals. A, anterior; D, dorsal. 3dpf.

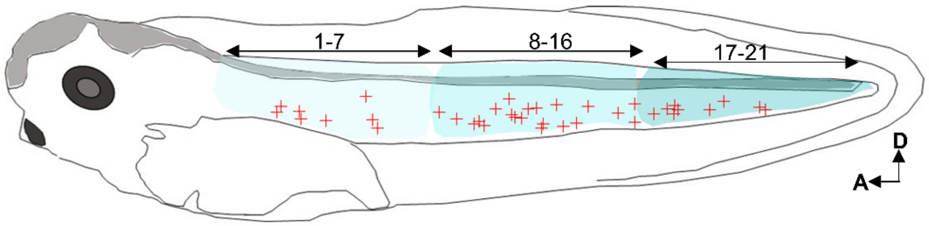

**Suppl Fig 8. A-P axis distribution of motor neuron axon terminals in $GABA_ARαβγ$ larvae that express GABA.** The distribution of GABA+ axon terminals (+) from 20 different 3dpf $GABA_ARαβγ$ larvae are overlaid on the schematic. Each chevron (1-21) along the A-P axis was systematically examined and the positions of GABA axon terminals were marked. Light blue, chevrons 1 to 7; blue, chevrons 8-16; dark blue, chevrons 17-21. A, anterior; D, dorsal.

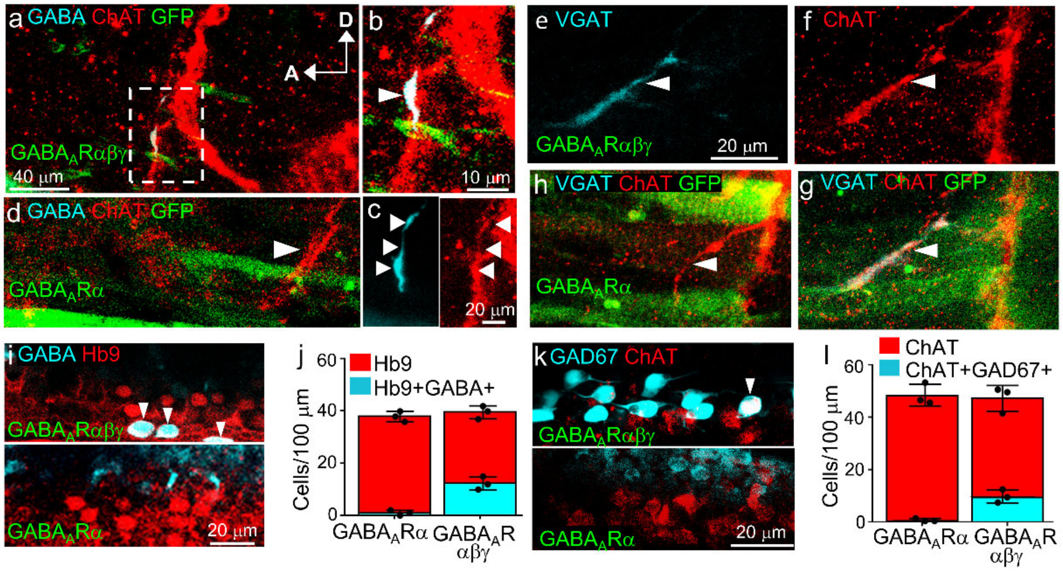

**Suppl Fig 9. GABAergic markers are expressed in motor neurons contacting GABA$_A$Rαβγ–GFP–expressing myocytes. (a)** GABA$_A$Rαβγ (green) myocyte is contacted by a GABA+ChAT+ axon. **(b)** Higher magnification of boxed region in a. **(c)** Individual channels for GABA+ and ChAT+ axon shown in b. **(d)** GABA is not detected in ChAT+ axons contacting a GABA$_A$Rα (green) myocyte. A, anterior, and D, dorsal, for all panels. **(e-g)** GABA$_A$Rαβγ myocyte is contacted by a VGAT+ChAT+ axon. **(h)** VGAT is not detected in ChAT+ axons contacting a GABA$_A$Rα myocyte. **(i)** *Top.* When myocytes express GABA$_A$Rαβγ, Hb9+GABA+ somas (arrowheads) are observed in the spinal cord. *Bottom*. Colocalization of Hb9 and GABA is not detected when myocytes express GABA$_A$Rα. **(j)** Quantification of cells per 100 μm of spinal cord (SC). **(k)** *Top, w*hen myocytes express GABA$_A$Rαβγ, a ChAT+GAD67+ soma (arrowhead) is observed in the SC. *Bottom,* colocalization of ChAT and GAD67 is not detected when myocytes express GABA$_A$Rα. **(l)** Quantification of (k). Error bars, S.D. All data from 3dpf larvae.

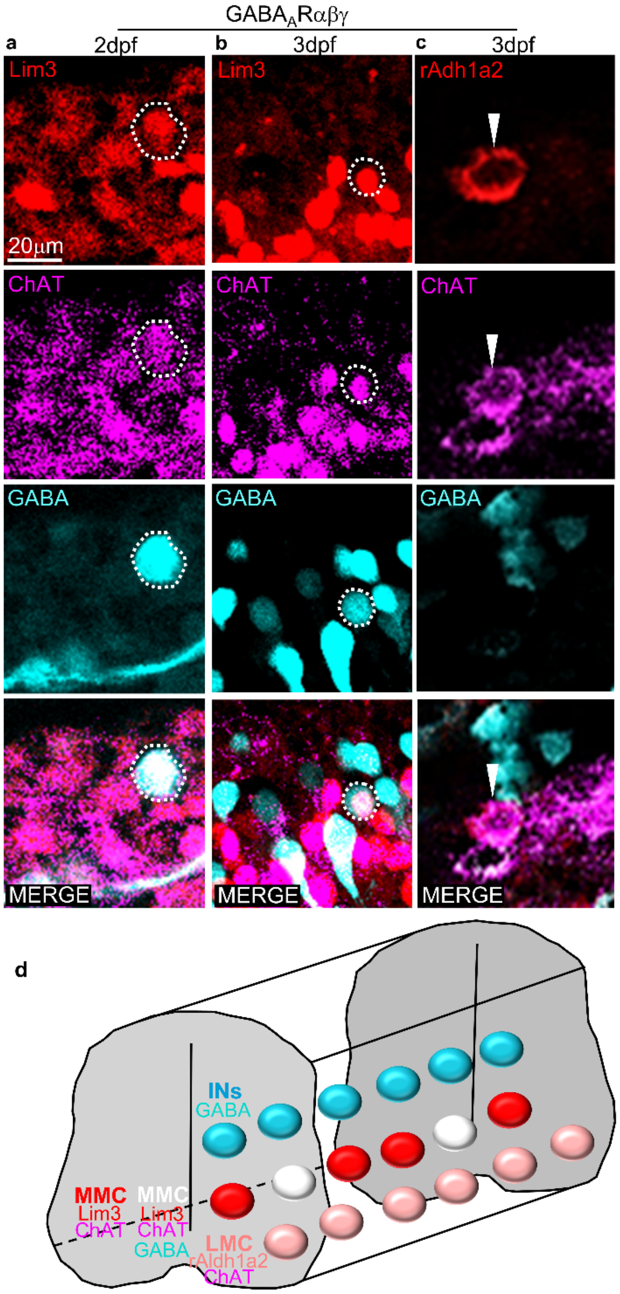

**Suppl Fig 10. ChAT+GABA+ motor neurons are derived from medial motor column motor neurons in GABA$_A$Rαβγ–expressing larvae.** Larvae injected with GABA$_A$Rαβγ mRNA in ventral blastomeres (V) were immunostained for medial motor column (MMC) marker Lim3, ChAT and GABA at 2dpf (**a**) and 3dpf (**b**). Dashed circles show Lim3+ChAT+GABA+ cells in the spinal cord. (**c**) Lateral motor column (LMC) marker rAdlh1a2, absent at 2dpf and detected at 3dpf, is positive only for ChAT (arrowhead) but not GABA. (**d**) Schematic shows the topographical location of ChAT+GABA+ (white) motor neurons in the MMC of the spinal cord. INs, interneurons.

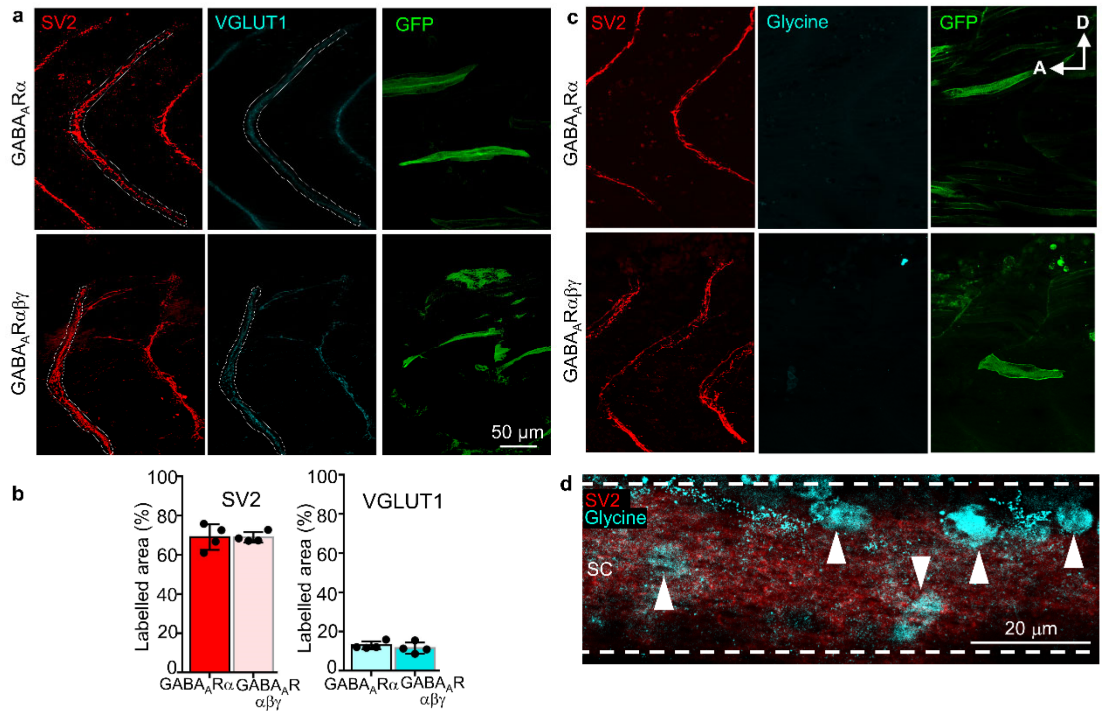

**Suppl Fig 11. Expression of GABA$_A$Rαβγ in ventral blastomeres did not produce detectable changes in expression of VGLUT1 or glycine in innervating axons.** (**a**) Embryo injected at the 4-cell stage with GABA$_A$Rα or GABA$_A$Rαβγ mRNA in the ventral blastomeres (V) and immunostained for SV2, VGLUT1 and GFP at 3dpf. Dotted lines outline the myocommatal junctions analyzed for SV2 and VGLUT1 area quantified in (**b**). (**c**) Embryo injected at 4-cell stage with GABA$_A$Rα or GABA$_A$Rαβγ mRNA in the V blastomeres and immunostained for SV2, glycine and GFP at 3dpf. (**d**) Example of glycine+ cell bodies (arrowheads) in the spinal cord (SC) of GABA$_A$Rα larva at 3dpf. A, anterior; D, dorsal.

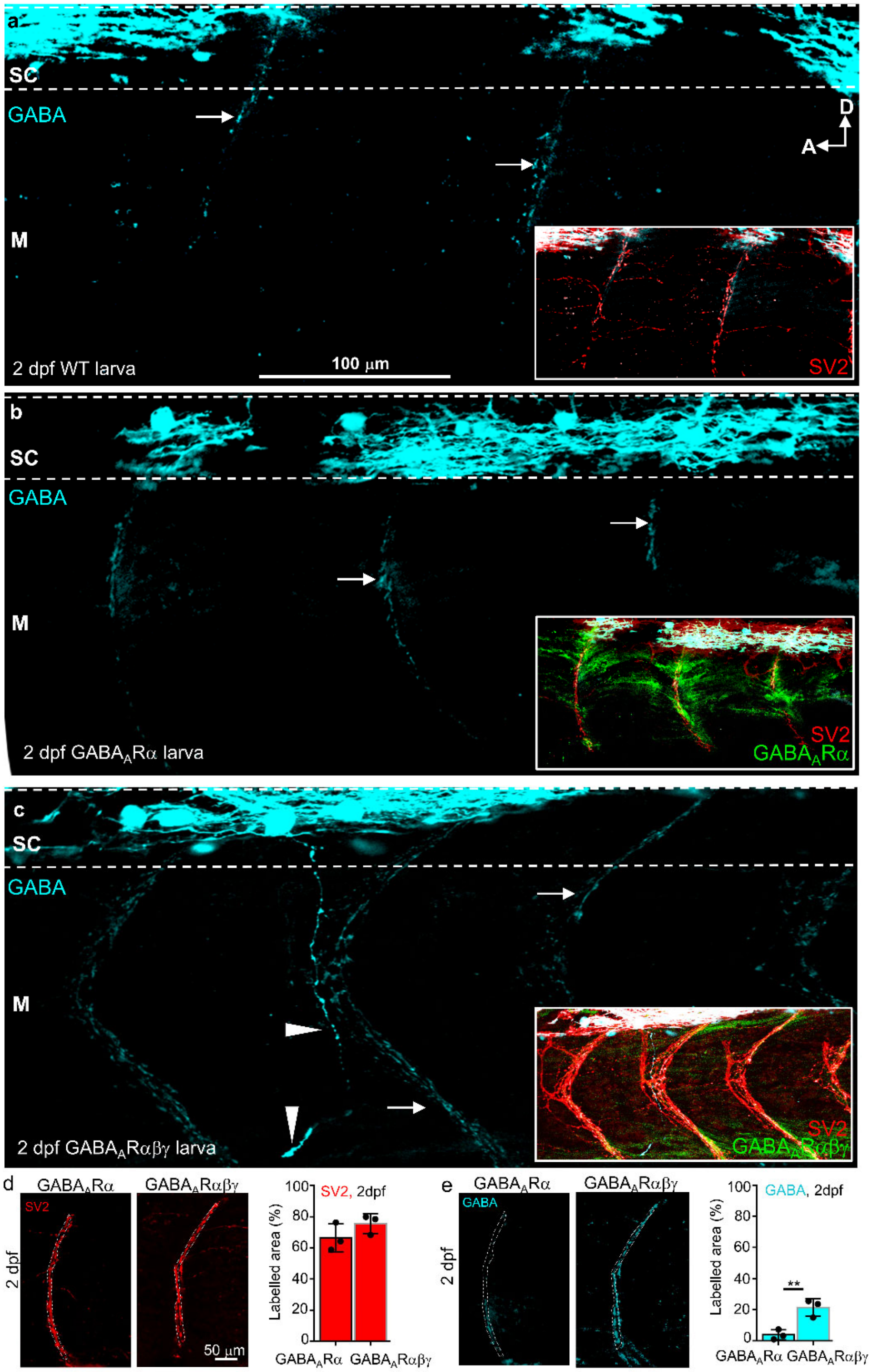

**Suppl Fig 12. GABA$_A$Rαβγ expression in myocytes stabilizes GABA expression in axons during development.** 2 dpf axonal GABA expression in the myotome of a wild type larva (**a**) and GABA$_A$Rα larva (**b**) has declined compared to 1dpf (cf Sfig. 6). (**c**) 2dpf GABA expression in the myotome of a GABA$_A$Rαβγ larva has been stabilized. Insets show SV2 and GABA$_A$R expression. A, anterior; D, dorsal. SC, spinal cord.

(**d, e**) Quantification of 2 dpf expression of SV2 and GABA. Dotted lines illustrate regions of myocommatal junctions analyzed for quantification of staining intensities. n=3 larvae, **p 0.0093.

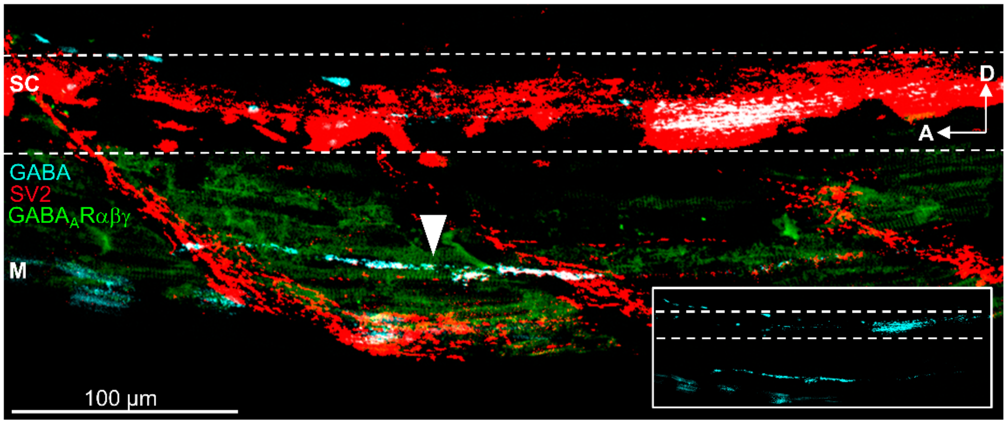

**Suppl Fig 13. GABA+ processes persist in the myotome of 7 dpf GABA$_A$Rαβγ larva.** GABA+SV2+ axon terminal contacting GABA$_A$Rαβγ (GFP+) myocytes (arrowhead). *Inset,* GABA+ axons in the myotome and spinal cord. A, anterior; D, dorsal. SC, spinal cord. M, myotome.

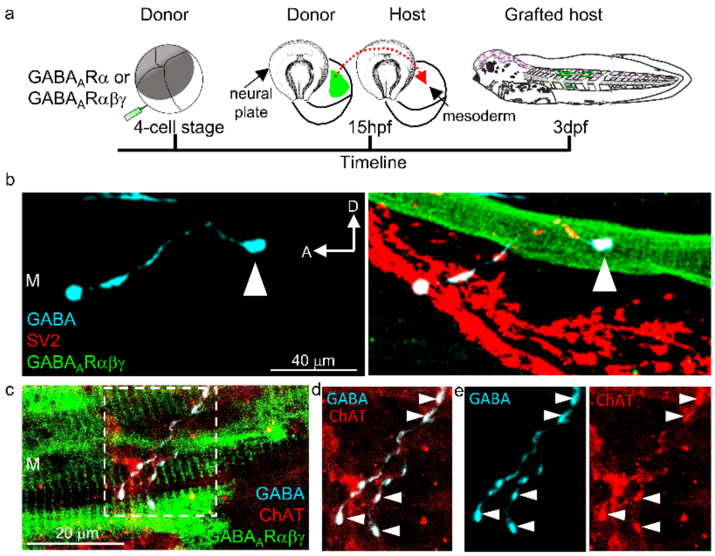

**Suppl Fig 14. GFP-labelled mesoderm grafts demonstrate the specificity of target tissue for $GABA_A$Rαβγ-dependent GABA expression in axon terminals.** (**a**) Schematic of mesodermal transplantation. *Left to right:* donor *Xenopus* embryo injected at the 4-cell stage with $GABA_A$Rα or $GABA_A$Rαβγ mRNA in the ventral blastomeres (V). Neural plate is lifted to access the presomitic mesoderm (green), which is grafted from donor to wild-type host larva at 15 hpf (red dashed arrow). GFP expression is detected in the myotome of host larva at 3dpf. (**b**) *Left:* GABA+ axons (cyan) course ventrally and posteriorly over the trunk myotome (M) in $GABA_A$Rαβγ–expressing-mesoderm-grafted host. *Right*: A GABA+SV2+ axon (arrowhead) contacts a $GABA_A$Rαβγ–expressing GFP+ myocyte. (**c**) Another $GABA_A$Rαβγ (green) myocyte is contacted by a GABA+ChAT+ axon. (**d**) Higher magnification of boxed region in c. (**e**) Individual channels for GABA+ and ChAT+ axon shown in d. 3dpf. D, dorsal. A, anterior.

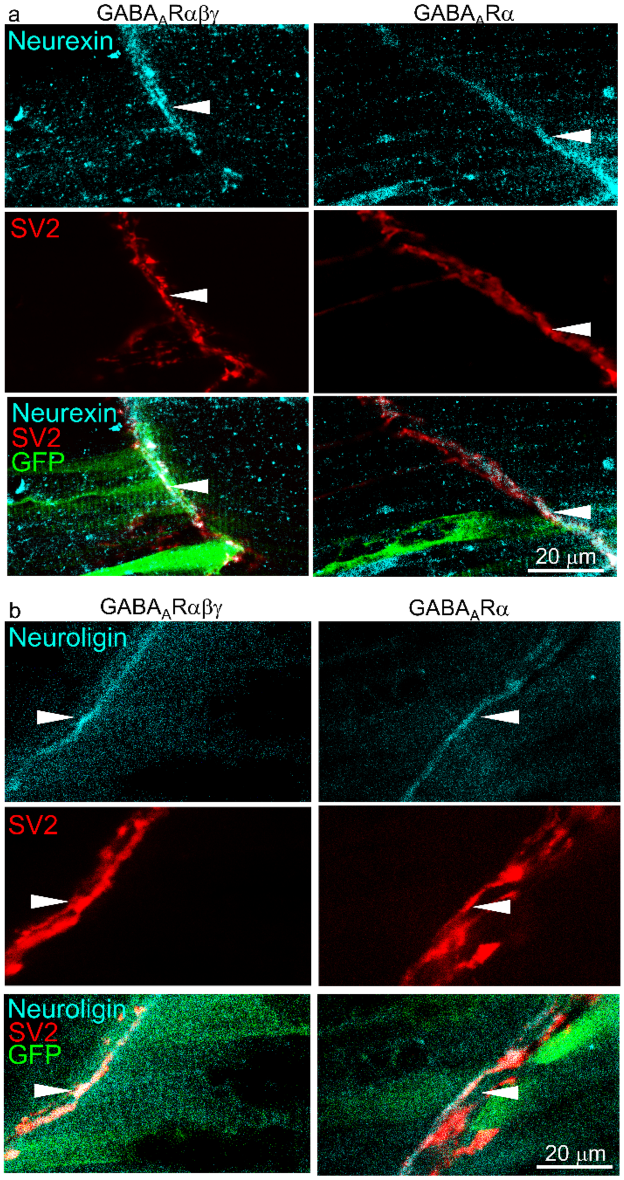

**Suppl Fig 15. Neurexin and neuroligin expression at the *Xenopus laevis* NMJ.** Neurexin (**a**) and neuroligin (**b**) expression is detected along SV2-labelled myocommatal junctions (arrowheads) of larvae expressing $GABA_AR\alpha\beta\gamma$ (left panels) and $GABA_AR\alpha$ (right panels). 3dpf.

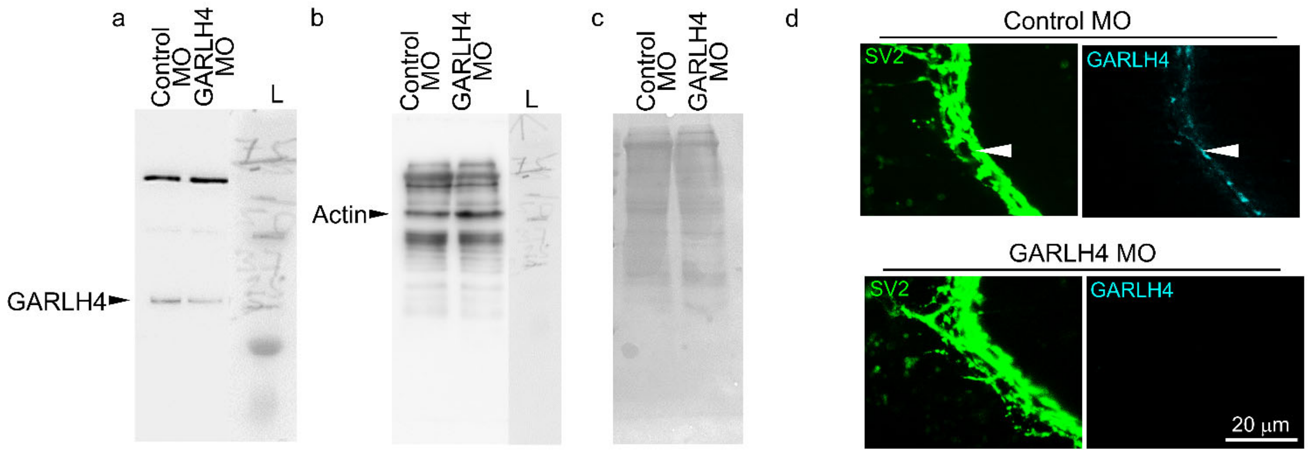

**Suppl Fig 16. Validation of the presence of GARLH4 in the *Xenopus* myotome and its knockdown by a GARLH4 MO.** $GABA_AR\alpha\beta\gamma$–expressing larvae were injected with a control MO or a GARLH4 MO (3 nl of 1 mM MO) in ventral blastomeres (V). Myotome protein lysates of these larvae were probed by Western blot for expression of GARLH4 (**a**) and the same blot was re-probed for the loading control, actin (**b**). (**c**) Total protein loading control with Ponceau S staining. Arrowheads indicate the positions of the respective proteins. L, molecular weight ladder. (**d**) GARLH4 expression is detected along SV2-labelled myocommatal junctions (arrowheads) of larvae expressing the control MO (top panels) but not in GARLH4 MO-expressing larvae (bottom panels). 3dpf.

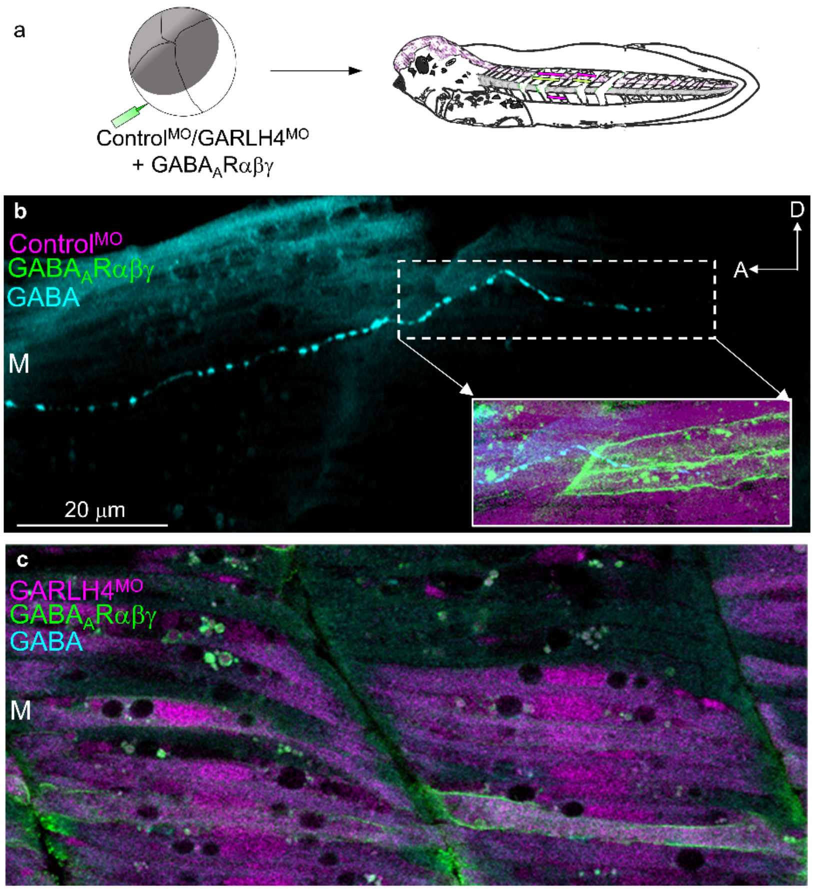

**Suppl Fig 17. Postsynaptic GABA$_A$Rαβγ plus GARLH4$^{MO}$ prevents induction of presynaptic GABA.** (**a**) Simultaneous injection of ventral blastomeres (V) with GABA$_A$Rαβγ mRNA along with control morpholino (control$^{MO}$) or GARLH4$^{MO}$. (**b**) GABA+ axon observed in myotome of control$^{MO}$ GABA$_A$Rαβγ larva. *Inset,* GABA+ axon contacts a GFP+ myocyte (7/7). (**c**) No GABAergic process is evident contacting myocytes expressing GARLH4$^{MO}$ and GABA$_A$Rαβγ (0/12). M, myotome. A, anterior; D, dorsal. n/N, larvae with GABA+ axon/total larvae examined.

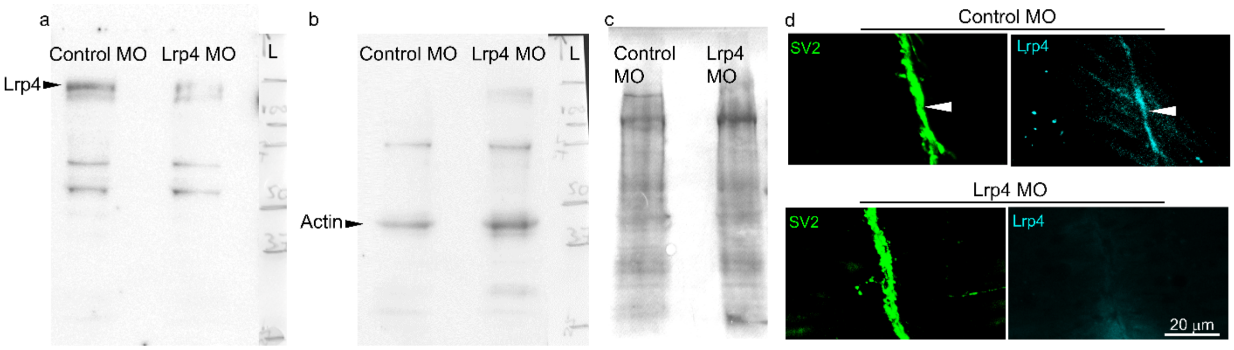

**Suppl Fig 18. Validation of the presence of Lrp4 in the *Xenopus* myotome and its knockdown by a Lrp4 MO.** Wild type larvae were injected with control morpholino (MO) or Lrp4 MO (6 nl of 1 mM MO) in ventral blastomeres (V). Myotome protein lysates of these larvae were probed by Western blot for expression of Lrp4 (**a**) and the same blot was re-probed for the loading control, actin (**b**). (**c**) Total protein loading control with Ponceau S staining. Arrowheads indicate the positions of the respective proteins. L, molecular weight ladder. (**d**) Lrp4 expression is detected in SV2-labelled myocommatal junctions (arrowheads) of larvae expressing the control MO (top panels) but not in Lrp4 MO-expressing larvae (bottom panels). 3dpf.

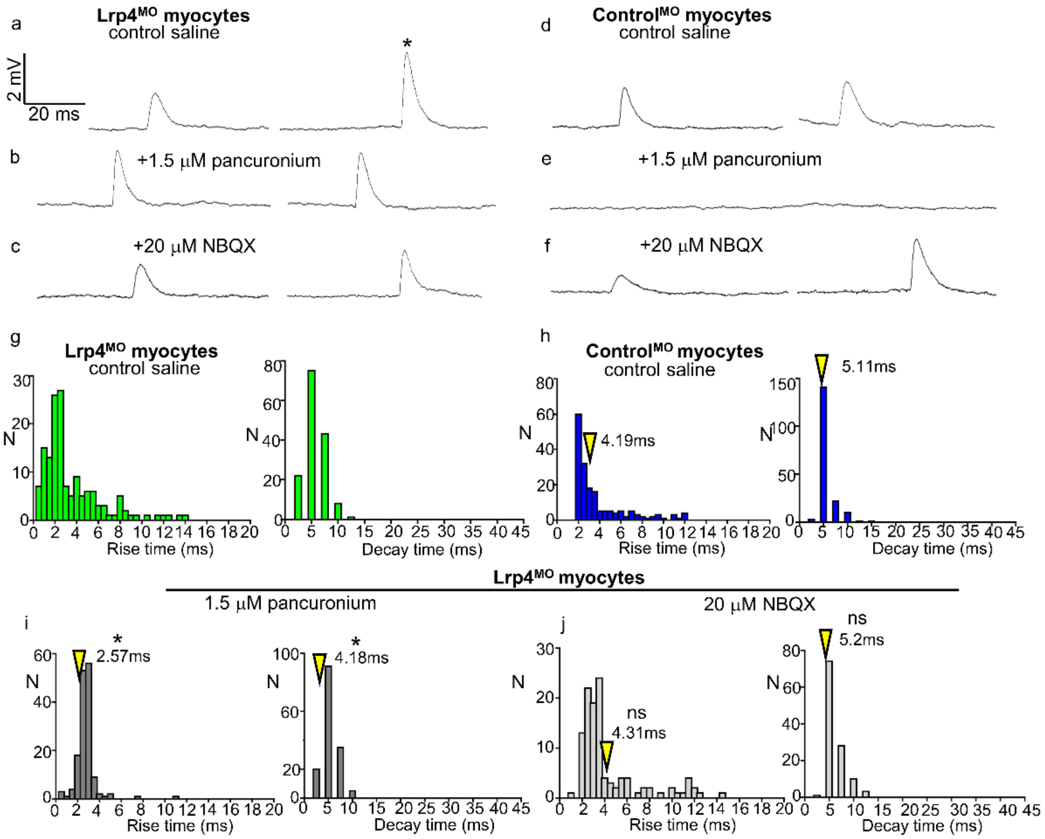

**Suppl Fig 19. Neuromuscular junctions of myocytes that express the Lrp4 morpholino generate glutamatergic and cholinergic mEPPs.** (**a-c**) Recordings from Lrp4$^{MO}$-expressing myocytes of 4dpf larvae reveal mEPPs with rise and decay times similar to AMPAR PSPs (asterisk) that are pancuronium-resistant and NBQX-sensitive, as well as mEPPs with rise and decay times similar to those described for nicotinic receptor mediated mEPPs that are pancuronium-sensitive and NBQX-resistant. (**d-f**) Recordings from control$^{MO}$-expressing myocytes reveal only pancuronium-sensitive mEPPs with rise and decay times similar to those for nicotinic receptor mediated mEPPs. (**g,h**) Rise and decay time distributions for mEPPs in myocytes of Lrp4$^{MO}$ larvae and control$^{MO}$ larvae in presence of saline. (**i,j**) Rise and decay time distributions for mEPPs in myocytes of Lrp4$^{MO}$ larvae in presence of pancuronium or NBQX. N, number of mEPPs. $\geq$ 155 mEPPs ($\geq$7 tadpoles) for each group. Only mEPPs with decay times fit by single exponentials were included. Resting potentials were held near -60 mV. Arrowheads indicate median values. Kolmogorov-Smirnov test compared rise time and decay time in (**h**) with respective rise and decay time in (**i**) and (**j**). *p<0.05, ns not significant.

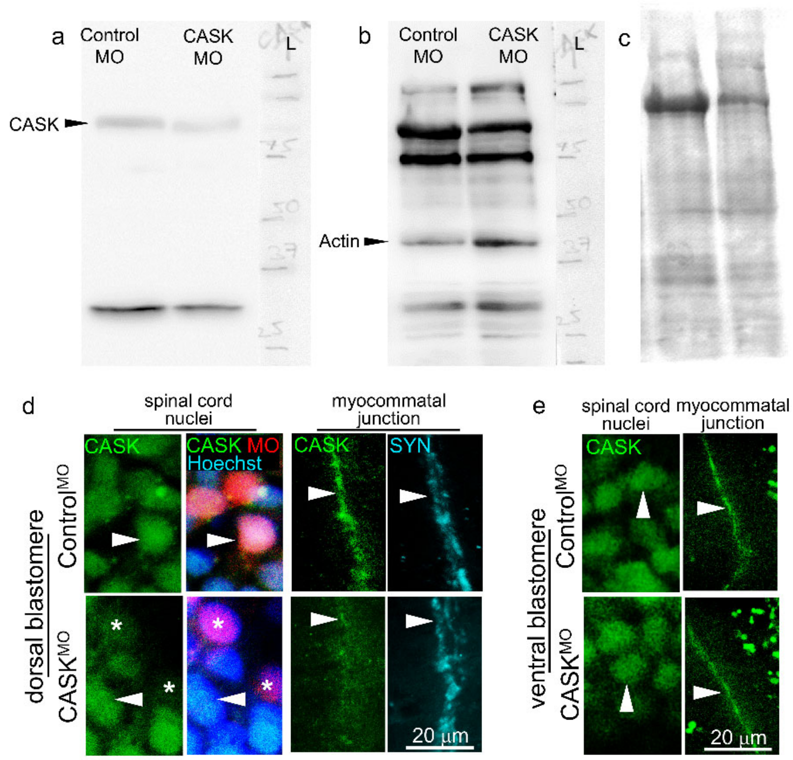

**Suppl Fig 20. Validation of presence of CASK in the *Xenopus* spinal cord and its knockdown by a CASK MO.** Wild type larvae were injected with control MO or CASK MO (6 nl of 1 mM MO) in dorsal blastomeres (D1.2) at 16-cell stage. Spinal cord protein lysates of these larvae were probed by Western blot for expression of CASK (**a**) and the same blot re-probed for the loading control, actin (**b**). (**c**) Total protein loading control with Ponceau S staining. Arrowheads indicate the positions of the respective proteins. L, molecular weight ladder. (**d**) CASK expression was detected in the Hoechst-labelled nuclei of cell bodies in spinal cord (arrowheads) of larvae expressing the control MO (top panels) and reduced in CASK MO-expressing larvae (bottom panels: asterisk, CASK MO+ cells; arrowhead, CASK MO- cells). CASK expression was detected along synaptophysin (SYN)-labelled myocommatal junctions (arrowhead) and reduced upon CASK knockdown. (**e**) No change was detected in expression of CASK in the myocommatal junctions and in spinal cord cell bodies following knockdown of CASK in myocytes. 3dpf.

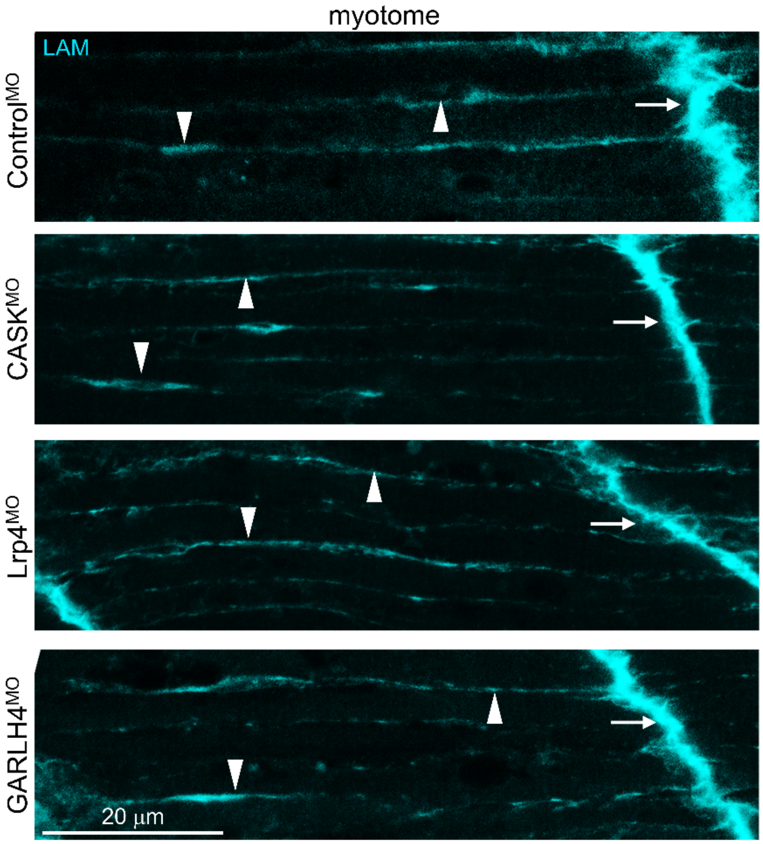

**Suppl Fig 21. No detectable change in laminin expression following morpholino injections.** Wild type larvae were injected with control$^{MO}$ or CASK$^{MO}$ in dorsal blastomeres (D1.2) at 16-cell stage or Lrp4$^{MO}$ or GARLH4$^{MO}$ in ventral blastomeres (V) at 4 cell stage. No change was detected in laminin (LAM) expression in MO-expressing larvae along the myocytes (arrowheads) and in the myocommatal junctions (arrows). 6 nl of 1 mM MO injected. 3dpf.

| ***Bead implanted*** | ***Larvae imaged (N)*** | ***Immuno for*** | ***Observation*** | ***P value (Control vs Treatment)*** |
|---|---|---|---|---|
| ***3 dpf (St 41)*** | | | **Larvae with altered (reduced) ChAT in myocommatal junctions** | |
| Control (Saline) | 5 | ChAT, SV2 | 0 | **0.0025 |
| Pancuronium | 13 | ChAT, SV2 | 11 | |
| ***4 dpf (St 43)*** | | | **Larvae with altered (reduced) ChAT in myocommatal junctions** | |
| Control (Saline) | 13 | ChAT, SV2 | 0 | ***0.0001 |
| Pancuronium | 19 | ChAT, SV2 | 13 | |
| | | | **Larvae with altered NT marker (increased, vGluT1/GABA/Glycine) in myocommatal junctions** | |
| Control (Saline) | 8 | vGluT1, SV2 | 0 | **0.0017 |
| Pancuronium | 14 | vGluT1, SV2 | 10 | |
| Control (Saline) | 5 | GABA, SV2 | 0 | |
| Pancuronium | 5 | GABA, SV2 | 0 | |
| | | | **Larvae with altered NTR marker (increased GluR1, NR1) in chevrons** | |
| Control (Saline) | 6 | GluR1, SV2 | 0 | ***0.0005 |
| Pancuronium | 15 | GluR1, SV2 | 13 | |
| Control (Saline) | 8 | NR1, SV2 | 0 | **0.0040 |
| Pancuronium | 10 | NR1, SV2 | 7 | |
| | | | **Larvae with altered (reduced) ChAT in myocommatal junctions** | |
| Control (Saline) | 8 | ChAT, SV2 | 0 | ***0.0007 |
| dTC | 11 | ChAT, SV2 | 9 | |
| ***2 dpf (St 36)*** | | | **Larvae with altered (reduced) ChAT in myocommatal junctions** | |
| Control (Saline) | 7 | ChAT, SV2 | 0 | ***0.0001 |
| Pancuronium | 18 | ChAT, SV2 | 15 | |
| | | | **Larvae with altered NT marker (increased vGluT1/Glycine) in myocommatal junctions** | |
| Control (Saline) | 8 | vGluT1, SV2 | 0 | **0.0010 |
| Pancuronium | 14 | vGluT1, SV2 | 11 | |
| Control (Saline) | 5 | GABA, SV2 | 0 | |
| Pancuronium | 5 | GABA, SV2 | 0 | |
| ***Morpholino experiments 3dpf (St 41)*** | | | **Larvae with altered (reduced) ChAT in myocommatal junctions** | |
| Lrp4 inverted morpholino (control) | 4 | ChAT, SV2 | 0 | 0.1429 |
| Lrp4 morpholino | 4 | ChAT, SV2 | 3 | |
| Lrp4 inverted morpholino (control) | 5 | ChAT, SV2 | 0 | **0.0047 |
| Lrp4 morpholino (3x increased conc) | 8 | ChAT, SV2 | 7 | |
| CASK inverted morpholino (control) | 4 | ChAT, SV2 | 0 | |
| CASK morpholino | 12 | ChAT, SV2 | 9 | *0.0192 |
| | | | **Larvae with altered (reduced) CASK in myocommatal junctions** | |
| Control morpholino (vendor supplied) | 4 | CASK, Synaptophysin | one-way ANOVA comparing Control MO, Lrp4MO, $GABA_A R\alpha\beta\gamma$+Lrp4MO, $GABA_A R\alpha\beta\gamma$+GARLH4MO conditions | $F_{3,12}$=6.316; p=0.0081; ControlMO vs Lrp4MO=*0.0109 |
| Lrp4 morpholino (3x increased conc) | 4 | CASK, Synaptophysin | | |
| | | | **Larvae with altered (reduced) CASK in spinal cord nuclei** | |
| Control morpholino (vendor supplied) | 5 | CASK, Hoechst | one-way ANOVA comparing Control MO, Lrp4MO, $GABA_A R\alpha\beta\gamma$+Lrp4MO, $GABA_A R\alpha\beta\gamma$+GARLH4MO conditions | $F_{3,12}$=16.5; p<0.0001; ControlMO vs Lrp4MO=ns |
| Lrp4 morpholino (3x increased conc) | 5 | CASK, Hoechst | | |

**Suppl Table 1 Loss of Function experiments.** Control and treatment data compared using Fisher's exact test (unless otherwise stated) for calculating percentage success - change in neurotransmitter expression along myocommatal junctions in each group.

| *Receptor overexpressed* | *Larvae imaged (N)* | *Immuno for* | *Observation* | *P value (GABA$_A$R αβγ vs GABA$_A$R α)* |
|---|---|---|---|---|
| ***3 dpf (St 41)*** | | | **Larvae with GABA in peripheral axons** | |
| GABA$_A$Rαβγ | 13 | GABA, SV2, GFP | 12 | ** 0.0071 |
| GABA$_A$Rα | 3 | GABA, SV2, GFP | 0 | |
| no injection | 3 | GABA, SV2, GFP | 0 | |
| | | | **Larvae with GABA in peripheral axons** | |
| GABA$_A$Rαβγ | 8 | GABA, ChAT, GFP | 8 | |
| GABA$_A$Rα | 5 | GABA, ChAT, GFP | 0 | ***0.0008 |
| no injection | 1 | GABA, ChAT, GFP | 0 | |
| GABA$_A$Rαβ | 9 | GABA, ChAT, GFP | 1 | |
| | | | **Larvae with GABA in peripheral axons** | |
| GABA$_A$Rαβγ | 14 | Hb9, GABA, GFP | 14 | |
| GABA$_A$Rα | 7 | Hb9, GABA, GFP | 0 | ****<0.0001 |
| | | | **Larvae with VGAT in peripheral axons** | |
| GABA$_A$Rαβγ | 6 | VGAT, ChAT, GFP | 6 | |
| GABA$_A$Rα | 6 | VGAT,ChAT, GFP | 0 | **0.0022 |
| no injection | 6 | VGAT | 0 | |
| | | | **Larvae with GABA in peripheral axons** | |
| GABA$_A$Rαβγ mesoderm transplantation | 5 | GABA, SV2, GFP | 5 | **0.0079 |
| GABA$_A$Rα mesoderm transplantation | 5 | GABA, SV2, GFP | 0 | |
| no transplantation | 3 | GABA, SV2, GFP | 0 | |
| | | | **Larvae with GABA in peripheral axons** | |
| ***1 dpf (St 22) WT*** | 8 | GABA, SV2 | 8 | |
| ***3 dpf (St 41)*** | | | **Larvae with altered NT marker (increased vGluT1/Glycine) in myocommatal junctions** | |
| GABA$_A$Rα | 4 | vGluT1, SV2 | 0 | |
| GABA$_A$Rαβγ | 4 | vGluT1, SV2 | 0 | |
| GABA$_A$Rα | 5 | Glycine, SV2 | 0 | |
| | 5 | Glycine, SV2 | 0 | |
| | | | **Larvae with GABA in peripheral axons** | |
| ***2 dpf (St 36) WT*** | 8 | GABA, SV2 | 6 | |
| ***7 dpf (St 48)*** | | | | |
| GABAARαβγ | 5 | GABA, SV2, GFP | 5 | **0.0079 |
| GABAARα | 4 | GABA, SV2, GFP | 0 | |
| ***Morpholino experiments 3dpf (St 41)*** | | | **Larvae with GABA in peripheral axons** | |
| GABA$_A$Rαβγ + GARLH4 (a) morpholino | 7 | GABA, Lissamine, GFP | 0 | ***0.0006 |
| GABA$_A$Rαβγ + Standard morpholino | 6 | GABA, Lissamine, GFP | 6 | |
| GABA$_A$Rαβγ + GARLH4 (c) morpholino | 8 | GABA, Lissamine, GFP | 0 | ***0.0006 |
| GABA$_A$Rαβγ + GARLH4 (c) morpholino (reduced concentration)- separate injections | 5 | GABA, Lissamine, GFP | 4 | ns |
| GABA$_A$Rαβγ + CASK morpholino (dorsal blastomere) | 4 | GABA, Lissamine, GFP | 1 | *0.0152 |
| GABA$_A$Rαβγ + CASK morpholino (dorsal blastomere, increased concentration) | 12 | GABA, Lissamine, GFP | 1 | ***0.0003 |
| | | | **Larvae with altered (reduced) CASK in myocommatal junctions** | |
| Control morpholino (vendor supplied) | 4 | CASK, Synaptophysin | one-way ANOVA comparing Control MO, Lrp4MO, GABA$_A$Rαβγ+Lrp4MO, GABA$_A$Rαβγ+GARLH4MO conditions | $F_{3,12}$=6.316; p=0.0081; ControlMO vs GABA$_A$Rαβγ+Lrp4MO=ns; Control MO vs GABA$_A$Rαβγ+GARLH4MO=ns |
| GABA$_A$Rαβγ+Lrp4 morpholino (3x increased conc) | 4 | CASK, Synaptophysin | | |
| GABA$_A$Rαβγ+GARLH4 (c) morpholino | 4 | CASK, Synaptophysin | | |
| | | | **Larvae with altered (reduced) CASK in spinal cord nuclei** | |
| Control morpholino (vendor supplied) | 5 | CASK, Hoechst | one-way ANOVA comparing Control MO, Lrp4MO, GABA$_A$Rαβγ+Lrp4MO, GABA$_A$Rαβγ+GARLH4MO conditions | $F_{3,12}$=16.5; p<0.0001; ControlMO vs GABA$_A$Rαβγ+Lrp4MO=***0.0001; Control MO vs GABA$_A$Rαβγ+GARLH4MO=ns |
| GABA$_A$Rαβγ+Lrp4 morpholino (3x increased conc) | 5 | CASK, Hoechst | | |
| GABA$_A$Rαβγ+GARLH4 (c) morpholino | 5 | CASK, Hoechst | | |

**Suppl Table 2 Gain of Function experiments.** GABA$_A$Rαβγ and GABA$_A$Rα data compared using Fisher's exact test (unless otherwise stated) for calculating percentage success - presence of GABA+ peripheral axons in each group.
